# Neutrophil Extracellular Trap Formation and Complement Activation Pathways Dominate Microbiota-dependent disease bias in lupus-prone female NZM2328 mice

**DOI:** 10.64898/2026.06.28.735075

**Authors:** Soumyabrata Roy, Jaisheela Vimali Irudhayaraj, Rekha Jalandra, Ping Lu, Diana-Christine Boucher, Radhika Gudi, Loni Carter, Caroline Westwater, Chenthamarakshan Vasu

**Author notes:** Address Correspondence: Chenthamarakshan Vasu, Department of Pharmacology and Immunology Medical University of South Carolina, 173 Ashley Avenue, BSB214B Charleston, SC-29425.

## Abstract

Women are predisposed to systemic lupus erythematosus (SLE) with a prevalence ratio of up to 9:1 over men. Multiple mouse strains including NZM2328 exhibit strong female dominance in developing spontaneous lupus as in humans with SLE. While lupus-prone mice can develop disease under germ free (GF) condition, the role of gut microbiota in female bias for lupus nephritis is not investigated systematically. Here, using specific pathogen free (SPF) and GF NZM2328 mice, and employing microbiota-depletion and microbial-association strategies, we show that microbiota influences lupus-like disease outcomes differently in males and females. Female NZM2328 mice with intact microbiota presents higher inflammation factor expression, including X-chromosome linked TLRs, in the distal gut and systemic compartments, and higher activation of genes and biological pathways such as neutrophil extracellular trap (NET) formation and complement and coagulation cascade (CCC) pathways, associating with their higher disease susceptibility. Gut microbiota-depletion as well as GF derivation eliminated not only the modest differences in the serum and fecal antibody levels and nAg reactivity, but also the gender bias in the timing of clinical stage disease onset as well as systemic NET and CCC pathway activation. Reciprocally, conventionalization of GF NZM2328 mice at juvenile age restored the female bias in intestinal and systemic autoantibody levels, pro-inflammatory immune pathway activation, and the timing of clinical stage disease onset. Overall, our observations show that, while genetic susceptibility appears to be the cause of lupus-like disease in NZM2328 mice, differential activation of NET and CCC pathways in males and females upon exposure to gut microbes, in combination with host-factors, causes gender bias in disease outcomes. We conclude that microbiota exposure-dependent protection of males and overactivation of NET and CCC pathways in females could be contributing to the female bias in lupus-like disease in NZM2328 mice.

## 1. Introduction

Systemic lupus erythematosus (SLE), an autoimmune disease, is caused by abnormally functioning B lymphocytes that produce autoantibodies to nuclear antigens including DNA and proteins [1]. These autoantibodies form immune complexes and cause damage to the tissues including kidney. A combination of genetic and multiple environmental factors contribute to the progression of disease in genetically susceptible subjects [1–3]. Many recent studies including ours have shown that gut microbiota modulates the rate of disease progression and the eventual overall disease outcome [4–8]. A key characteristic of SLE is the predisposition of women to develop this disease with a prevalence ratio of up to 9:1 over men [9–11] and studies using human samples and mouse models have shown a role for sex hormones in lupus-associated sex dimorphism [12–16]. By employing an F1 strain of lupus-prone (SWRxNZB)F1 (SNF1) mice, a spontaneous disease model with significant female bias, we have shown that microbiota-dependent pro-inflammatory immune response in the gut mucosa of females is initiated at juvenile ages as well as androgen-dependent protection of males at adult ages contributes to gender differences in the systemic autoimmune progression [17]. Our recent reports also showed that the abundance and nuclear antigen (nAg) reactivity of fecal IgA, which is largely gut microbiota-dependent, are predictive of the eventual systemic autoantibody levels and the disease-associated female bias in multiple strains of lupus-prone mice [18–20], and the systemic autoimmunity in SLE patients [21]. Overall, our previous studies suggested an involvement of gut microbiota in determining the degree of systemic autoimmunity and the female bias in disease progression/incidence under genetic susceptibility.

While it has been shown by others that lupus-prone mice do develop disease under germ-free (GF) condition and host genetics is the primary contributor to this disease process [22–24], it is unknown if female bias in lupus persists in relevant animal models after GF derivation. Studies using GF type 1 diabetes (T1D)-susceptible mice have shown that lack of exposure to gut microbiota results in the loss of gender bias in this T cell-mediated autoimmunity [25, 26]. Our report that used broad spectrum antibiotic treated as well as castrated SNF1 mice found that gut sex hormone - microbiota interactions determine the disease outcomes differently in males and females [17]. Nevertheless, it is not fully understood whether the dynamics of autoantibody production and disease progression are different in male and female lupus-prone mice under GF condition. In this regard, while F1 strains with lupus susceptibility (SNF1 and NZB/WF1) show strong female dominance in autoantibody production and clinical stage disease onset, these strains are not logistically ideal for gnotobiotic studies. NZM2328 mouse strain, derived from extended intercrosses of NZB/WF1 mice [27] and with well-recognized female bias similar to that of SNF1 and NZB/WF1 mice and human SLE [28], on the other hand, is ideal for studies under gnotobiotic conditions. Of note, unlike in SNF1 and NZB/WF1 mice, lupus-like disease-associated female bias in NZM2328 is pronounced primarily in terms of kidney disease but not autoantibody levels [19, 28], and the role of gut microbiota on its disease-associated sex dimorphism has not been investigated.

Here, we show that female NZM2328 mice with intact microbiota (specific-pathogen free; SPF mice) present higher inflammation marker expression in the distal gut and systemic compartments, and higher activation of genes and biological pathways including neutrophil extracellular trap (NET) formation and complement and coagulation cascades (CCC) pathways, which associate with early disease onset, as compared to their male counterparts. Gut microbiota-depletion as well as GF derivation eliminated not only the differences in serum and fecal antibody levels and nAg reactivity, but also the gender bias in the onset of clinical stage disease onset as well as systemic NET and CCC pathway activation. Reciprocally, conventionalization of GF NZM2328 mice restored the female bias in intestinal and systemic autoantibody levels, NET and CCC pathway gene expression, and clinical stage disease onset. Overall, these observations show that, while host genetic factors appear to be the cause of lupus-like disease in NZM2328 mice, the gut microbiota-host factor interaction contributes to differential activation of NET and CCC pathways in males and females and cause female bias in disease outcomes.

## 2. Materials and Methods

### Mice

NZM2328 mouse [28] breeders were provided by Dr. Shu Man Fu (University of Virginia). Breeding colony of these mice was established at the animal facility of Medical University of South Carolina (MUSC). Germ free (GF) NZM2328 mice were derived from this colony with the help of Taconic Biosciences service, and the breeding colony was established in the gnotobiotic facility at MUSC. GF status of mice is tested periodically by standard anaerobic and aerobic culture methods. All experimental protocols were approved by the Institutional Animal Care and Use Committee (IACUC) of MUSC. All methods in live animals were carried out in accordance with relevant guidelines and regulations of this committee and performed according to National Institutes of Health (NIH) Guidelines on Humane Care and Use of Laboratory Animals.

In some studies, SPF NZM2328 mice were given a cocktail of broad-spectrum antibiotics (Abx) in drinking water as described in our recent reports [17, 29], starting at 4 weeks of age to deplete the gut microbiota. Depletion of gut microbiota was confirmed by culture of the fecal pellet suspension on brain heart infusion (BHI) agar plates under aerobic and anaerobic conditions as described before [17, 29]. Fresh urine and feces, and tail vein blood samples were collected from individual SPF and GF mice at timely intervals. In some experiments, mice at different ages were euthanized and tissues were collected for immunological, genomic, and histopathological analyses.

### Proteinuria and severe disease

Urine samples were tested for protein levels by Bradford assay (BioRad) against bovine serum albumin standards as described before [17, 19, 30]. SPF mice that showed high proteinuria (>10mg/ml) for two consecutive weeks or GF mice that showed high proteinuria upon biweekly testing were considered to have severe nephritis and clinical stage disease. For GF mice, end-stage disease symptoms such as severe puffy appearance (whole body edema) or body condition score of 1 were also considered the experimental endpoint/severe nephritis and removed from the isolators and tested to confirm the kidney disease before euthanasia.

### ELISA

Levels of antibodies against nucleohistone and dsDNA in mouse sera were determined by ELISA as described in our recent reports [17–19]. Briefly, nucleohistone (Millipore-Sigma) or dsDNA from calf thymus (Millipore-Sigma) were coated as antigen (0.5 μg/well), overnight, onto ELISA plate wells. DNA binding solution (Thermo Scientific or in-house preparation) in PBS and carbonate buffer (pH 9.6) respectively, were used for coating the plates with dsDNA and nucleohistone. To determine the autoantibody titers, serial dilutions of the sera were made and IgM, IgG, IgA isotypes, and IgG1, IgG2a, IgG2b, IgG2c and IgG3 subtypes against these antigens were detected using biotin-conjugated respective anti-mouse immunoglobulin isotype/subtype antibodies, followed by HRP-labeled streptavidin (Southern Biotech). In some assays, a single, optimized dilution of the serum samples collected from multiple groups were tested side-by-side to determine the relative OD values. The abundance and dsDNA and NH reactivities of fecal IgA features were tested as detailed in our recent reports [18, 19]. Serum testosterone and 17β-estradiol (estrogen) levels were determined using ELISA and EIA kits from ALPCO and Enzo lifesciences respectively. Fecal calprotectin levels were determined using mouse S100A8/S100A9 heterodimer DuoSet ELISA (R&D systems) as described in our previous reports [20, 21].

### FITC-dextran assay and detection of circulating bacterial DNA levels

To determine gut permeability, mice were given FITC-dextran (4-kDa FITC-dextran; 5 mg/mouse for 4-week-old mice and 10 mg/mouse for 8-week-old mice) by oral gavage, euthanized after 2h and plasma samples were examined for FITC-dextran concentration using a fluorescence plate reader (Tecan) against control samples spiked with known concentration of FITC-dextran as described before [20, 31]. To determine the circulating bacterial DNA levels, whole blood collected from different groups of euthanized mice by cardiac puncture was subjected to qPCR assay using universal 16S rDNA-specific primer sets as described before [20, 31]. In some assays, total DNA was extracted from plasma samples as described before [20], pellets were solubilized in a volume that was proportionate to the starting material (plasma) volume and subjected to qPCR for detection of relative 16S rDNA levels.

### Quantitative PCR for detecting gene expression

Total RNA was extracted from comparable sized pieces of distal ileum, distal colon, or spleen using Trizol reagent (Invitrogen). cDNA was prepared from total RNA using oligo-dT containing first strand synthesis kit (Thermo) and PCR was performed using SYBR-green master-mix (BioRad) and target-specific custom-made primer sets as described in our previous reports [17, 31, 32]. A CFX96 real time PCR machine (BioRad) was used, and relative expression of each factor was calculated by the 2Δ-CT cycle threshold method against house-keeping gene (β-actin) control.

### Conventionalization of GF NZM2328 mice

Four-week-old GF NZM2328 male and female mice were housed in SPF facility with pooled “dirty bedding” materials from SPF NZM2328 breeding cages. Bedding material was replaced with fresh dirty bedding from the same breeding cages, every other day, thrice. Fresh fecal pellets from individual mice were collected after 15 days and subjected to anaerobic and aerobic culture to confirm colonization by the microbes. In some experiments, 4-week-old male or female GF NZM2328 mice were given a suspension of fresh fecal pellets from 16-week-old male or female SPF NZM2328 mice by oral gavage and housed on dirty bedding of the same SPF NZM2328 donor cages. These mice were examined for serum autoantibody levels at different time-points and monitored for proteinuria for up to 40 weeks of age.

### 16S rRNA gene sequencing and bacterial community profiling

Total DNA was prepared from the fecal pellets for bacterial community profiling as detailed in our previous reports [8, 17, 29, 33–35]. DNA was amplified by PCR using 16S rRNA gene-V3-V4 region targeted amplicon primer sets, and sequencing was performed using Illumina NextSeq 2000 (P1 300 PE Flowcell) with the help of NC State University Genomic Sciences Laboratory (Raleigh, NC, USA) or Molecular Research LP (Shallowater, TX). Sequencing reads were fed into the Metagenomics application of BaseSpace (Illumina) for performing taxonomic classification of 16S rRNA gene amplicon reads using an Illumina-curated version of the GreenGenes taxonomic database which provided raw classification output at multiple taxonomic levels. The OTUs were also normalized and used for metagenomes prediction of Kyoto Encyclopedia of Genes and Genomes (KEGG) orthologs by employing PICRUSt2 [36] downstream analysis feature of NIAID’s (Nephele https://nephele.niaid.nih.gov/). MicrobiomeAnalyst [37] and Statistical Analysis of Metagenomic Profile Package (STAMP) [38] applications were employed for statistical analysis and visualization of data.

### Bulk RNAseq analysis of gene expression

Bulk RNAseq assay was done with the help of Novogene Inc using polyA mRNA isolated from the total RNA preparations sent to this service provider as done before [32] (detailed in supplemental method). Raw sequencing counts data was then normalized and subjected to visualization and statistical analysis employing web-based integrated Differential Expression & Pathway analysis (iDEP) 2.01 platform (https://bioinformatics.sdstate.edu/idep/ [39] as described in supplemental method.

### Statistical analysis

Cumulative values of experiments conducted in multiple batches or samples collected from multiple experiments are used for some figure panels. Mean, SD, and statistical significance (*p-value*) were calculated using GraphPad Prism, Microsoft Excel, and/or online statistical applications. Wilcoxon signed-rank test was employed, unless specified, for values from *in vitro* and *ex vivo* assays. Specific methods employed upon statistical analysis and visualization of RNAseq and 16S rRNA gene amplicon sequencing data are mentioned in figure legends. Log-rank analysis was performed to compare the timing of lupus-like disease incidence (severe proteinuria onset) of the test group with that of respective control group. Fisher’s exact test was used for comparing nephritis severity in test vs. control groups. A *p* value of ≤0.05 was considered statistically significant.

## Results

### SPF NZM2328 shows gender specific differences in autoimmune/pro-inflammatory features

The *l*upus-prone mouse strain NZM2328, which was derived from extended intercrosses of an F1 strain (NZB/WF1), presents gender bias and slow-paced disease progression similar to that of SNF1 and NZB/WF1 mice and human SLE [19, 27, 28]. It has been reported that these NZM strains, particularly NZM2410 and NZM2328, have varying genetic contributions from NZB and NZW and display variable penetrance of lupus-like phenotypes [27]. In contrast to a well-studied NZM2410 strain, with limited female bias, NZM2328 mice develop autoantibodies and acute and severe chronic glomerulonephritis with profound female predominance, albeit lack of robust differences in total serum autoantibody levels [19, 28]. Since reports on the use of this mouse model is limited, first we have conducted systematic studies to determine if the NZM2328 mice housed in our SPF facility present gender specific differences in various features including the timing of proteinuria onset, levels of circulating antibody isotype and subclass, autoantibody titers, and the gene expression profile of spleen tissues. As show in **Fig. 1A**, severe proteinuria was detected in females as early as 20 weeks of age compared to males at about 24 weeks of age. Forty-week disease incidence, defined by the timing of high proteinuria, was about 80% for female and under 30% for males.

**Fig. 1:**
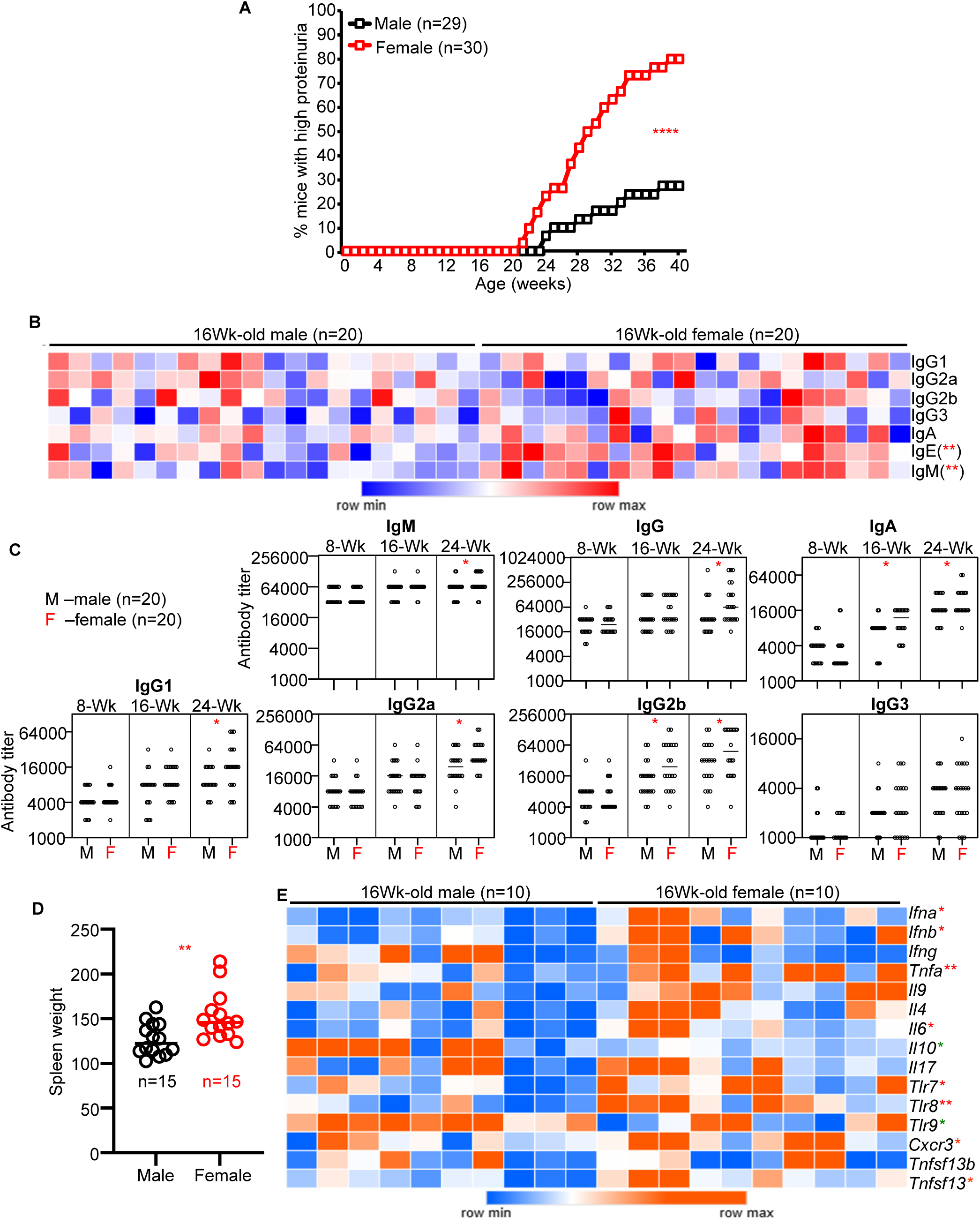
Gender specific differences in lupus-like disease and systemic immune features of SPF NZM2328 mice. **A)** Male and female NZM2328 mice housed in an SPF facility were tested for urine protein values every week for up to 40 weeks of age. Percentage of mice with severe nephritis as indicated by high proteinuria (≥10mg/ml) is shown. **B)** Serum samples collected at 16 weeks of ages were subjected to antibody isotyping/subtyping by ELISA (for IgA) or Luminex multiplex assay/suspension bead array (for the remaining isotypes/subtypes) and relative concentration values were used for generating the heatmap. **C)** Serum samples collected at different ages were subjected to ELISA to determine the dsDNA reactive titers of Ig isotypes and subclasses. NH reactive titers of the antibodies are shown in supplemental Fig. 1. **D)** Weight of spleens from mice euthanized at 16-weeks of age. **E)** RNA prepared from spleen tissues of 16-week-old mice were subjected to quantitative RT-PCR for the expression levels of various genes and the values relative to housekeeping gene (beta-actin) were used for generating the heatmap. *p-*values by log-rank test for panel A, Fisher exact test for panel B, Mann-Whitney test (comparison of males and females within specific age groups) for panels C, D, E and F. *<0.05, **0.01, ***0.001.

Quantification of serum antibody isotypes and IgG subtypes revealed that the total antibody concentrations were not significantly different in male and female NZM2328 mice at younger adult ages of up to 12 weeks (not shown), but the abundances of IgE and IgM were relatively higher in females at 16 weeks of age **(Fig. 1B)**. Serum nuclear antigen (nAg) reactive antibody (anti-dsDNA and anti-NH antibody) levels were determined in male and female NZM2328 mice at different ages. The overall IgA, IgM and IgG (as well as IgG isotype) titers against dsDNA and NH showed noticeable differences only at close to or at proteinuria onset stage (particularly at 16 and 24 weeks of age) in male and female mice **(Fig. 1C and supplemental Fig. 1).**

Although statistically significant, these differences appear to be modest relative to the difference in timing of severe proteinuria detection in males and females during the 40-week monitoring period. Like serum autoantibody levels, spleens of male and female showed weight heterogeneity with overall larger spleen mass in females compared to males **(Fig. 1D)** confirming splenomegaly in females. qPCR assay of cDNA prepared from whole spleen tissues revealed higher expression of many pro-inflammatory immune function-related genes such as *Ifna, Tgfb1, Tnfa, Il6, Tnfsf13* (APRIL), and X-chromosome-linked genes such as *Tlr7, Tlr8* and *Cxcr3* in females compared to their male counterparts at 16 weeks of age **(Fig. 1E).** On the other hand, *Il10* and *Tlr9* genes are expressed at relatively lower levels in female spleens compared to that of their male counterparts.

### NZM2328 mice show gender specific differences in the activation of multiple immune response-associated genes and biological pathways

Considering the subtle differences in systemic autoantibody levels between male and female NZM2328 mice, albeit profound gender bias in the 40-week disease incidence, bulk RNAseq assay was conducted using spleen tissues from male and female NZM2328 mice to identify differentially expressed genes and biological pathways. **Fig. 2A-C** show that several genes are differentially expressed in male and female NZM2328 mouse spleens at 16 weeks of age with females showing higher expression of a larger number of genes as compared to their male counterparts. Key gene expression overrepresented in females include neutrophil activation related genes such as *S100a8*, *S100a9*, *Mpo*, *Elane* and *Ngp*. On the other hand, expression of genes that encode for chaperone proteins such as *Hsph1*, *Hspa1a* and *Hspa1b* that manage protein folding and are involved in an unfolded protein response (UPR) are underrepresented in females compared to their male counterparts. Importantly, KEGG pathway enrichment analysis employing differentially expressed gene (DEG) profiles revealed an upregulation of many inflammatory and disease process-associated pathways, including those related to neutrophil extracellular trap (NET) formation, platelet activation, complement and coagulation cascades (CCC), and ECM-receptor interaction pathways in females compared to males **(Fig. 2D)**. On the other hand, protein processing in endoplasmic reticulum, antigen processing and presentation and estrogen signaling pathway function are significantly downregulated in the spleen tissues of female NZM2328 mice compared to males. Gene ontology (GO) enrichment analysis for biological processes employing DEG list also showed differential activation of similar pathways in female and male NZM2328 mouse spleens **(Supplemental Fig. 2)**. Overall, this gene expression profiling suggests that the systemic autoimmune processes in NZM2328 mice, females particularly, is facilitated by NET, CCC, platelet activation and ECM-receptor interaction pathway activation as well as defective protein folding and antigen processing functions.

**Fig. 2:**
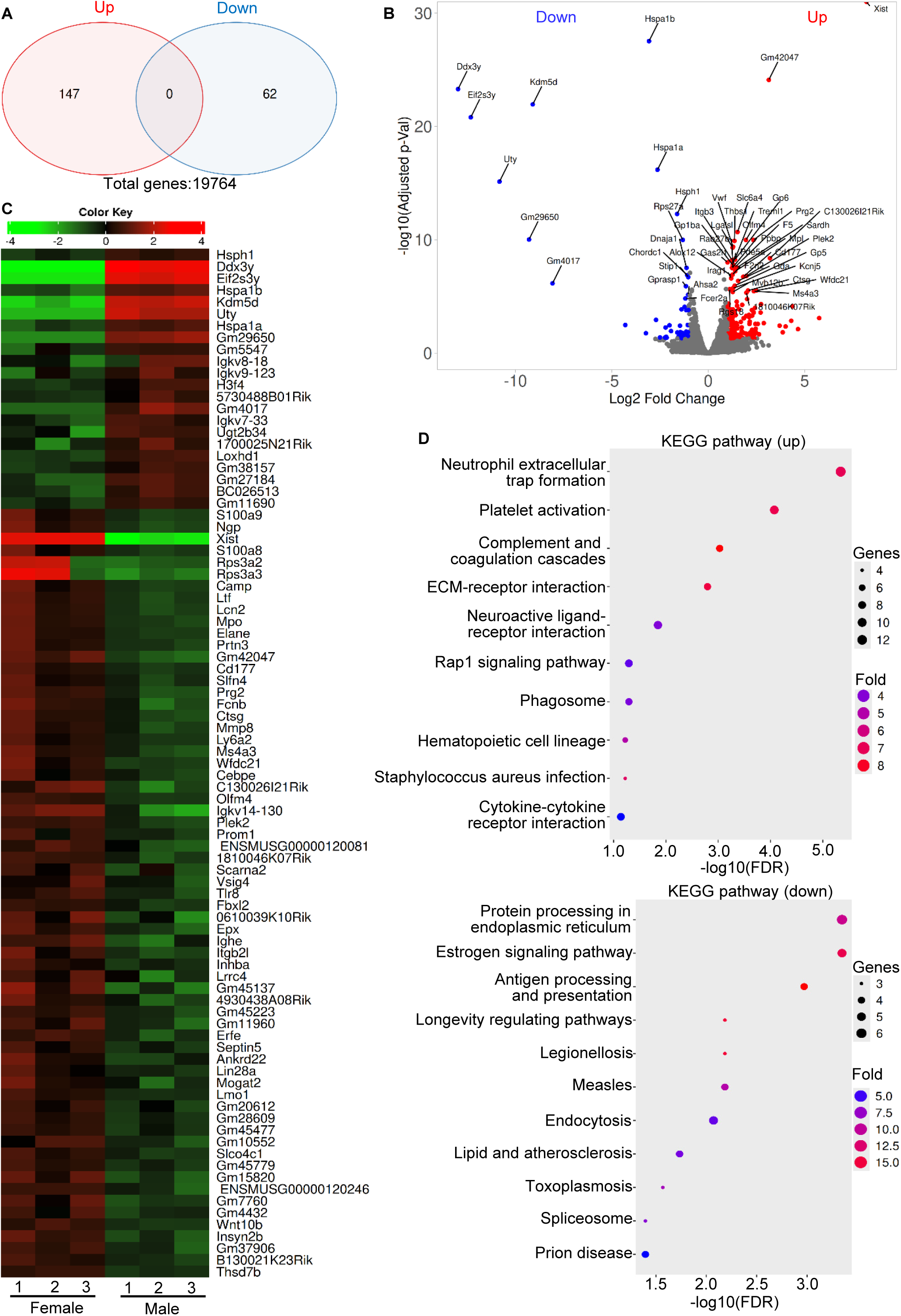
Differences in gene expression profiles of the spleen tissues from male and female NZM2328 littermates of SPF facility. RNA prepared from the spleen of 16-week-old male and female littermates of NZM2328 mice were subjected to bulk-RNAseq and the gene counts of data were then normalized and subjected to visualization and statistical analysis using iDEP2.01 platform. **A)** Venn diagrams showing number of significantly up- and down- regulated genes among differentially expressed genes (DEGs; identified employing DESeq2 with a minimum fold change of 2 and FDR cutoff as 0.05) in females compared to males. **B)** Volcano plot showing significantly up- and down- regulated genes among DEGs in females compared to males with top genes labeled. **C)** Heatmap showing top differentially expressed genes in females and males. D) KEGG Pathway enrichment analysis of significantly up- and down- regulated metabolic pathways or signal transduction pathways associated with DEGs in females compared to males. Gene ontology (GO) enrichment analysis of significantly up- and down- regulated biological processes in females is shown in supplemental Fig. S2.

### NZM2328 mice present gender specific differences in the gut inflammatory features and gene expression profiles

Our recent reports have shown a higher abundance and nAg reactivity of fecal IgA in lupus-prone mouse strains including NZM2328 as compared to B6 mice [18, 19]. However, unlike the hybrid SNF1 and NZB/W-F1 mouse models, NZM2328 mice did not show gender specific differences in fecal IgA features at juvenile or young adult ages [19]. Considering the female bias in lupus-like disease and gender specific differences in the activation of various inflammatory pathways in the systemic compartment, here we examined the fecal IgA features and calprotectin levels in a larger cohort of NZM2328 mice. As shown in Fig. 3A, while fecal IgA abundances were not significantly different, dsDNA and nAg reactivities of fecal IgA were considerably higher at 16 weeks of age. Further, Fig. 3B shows that fecal calprotectin levels are significantly higher in female mice as early as at 8 weeks of age as compared to their male counterparts. The degree of gut permeability (measured by FITC dextran method) or microbial translocation (based on serum bacterial DNA levels) were also relatively higher in females compared to males **(Fig. 3C&D)**. Examination of immune and gut epithelial barrier function gene expression by qPCR revealed relatively higher expression of some of the pro-inflammatory cytokine and X-chromosome-linked TLR genes and lower expression of tight junction associated *Ocln* and *Cldn-2* genes in the distal colon tissues of females **(Fig. 3E)**. As shown in our previous report which studied SNF1 mice [17], some of these differences were visible at as early as 4 weeks of age (not shown). However, differences in the expression levels of these genes in males and females were less pronounced in the distal ileum. Bulk RNAseq of distal colon tissue also revealed differential expression of large number of genes by male and female NZM2328 mice **(supplemental Fig. 3)**. KEGG pathway enrichment analysis of DEG in the distal colon tissue showed higher expression of multiple genes related to ECM-receptor interaction, AGE-RAGE (including PI3K-Akt) signaling and CCC pathways and modestly diminished expression of steroid biosynthesis and tight-junction function associated genes **(Fig. 3F)**. Overall, our results suggest that these pro-inflammatory features of distal gut may be contributing to the progression of systemic autoimmunity, and the higher disease incidence/early disease onset in female NZM2328 mice.

**Fig. 3:**
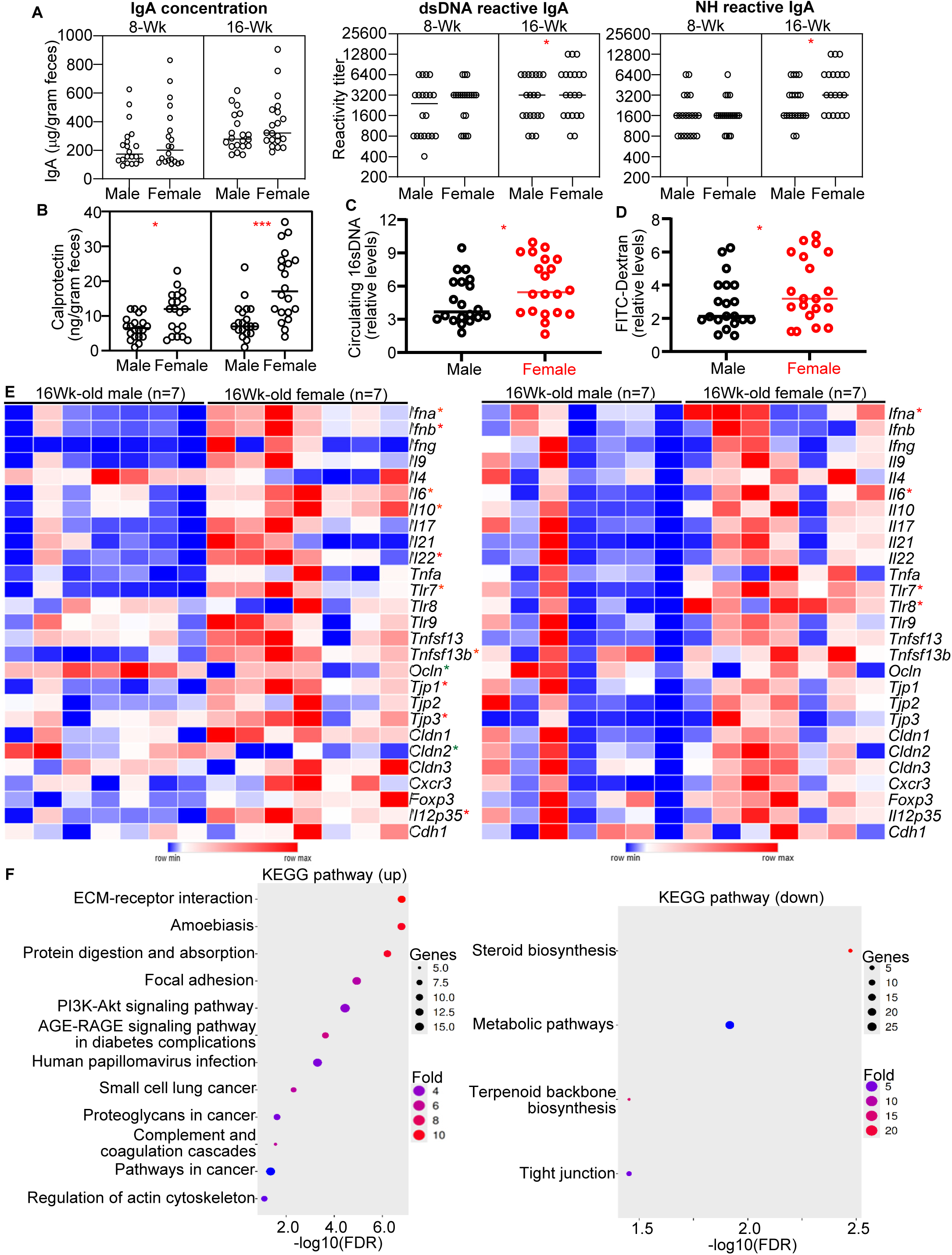
Gender specific differences in intestinal immune and permeability features and gene expression profiles in SPF NZM2328 mice. Extracts of fecal samples from different age groups of male and female SPF NZM2328 mice were examined for **A)** total IgA concentration and dsDNA and NH reactive IgA titers and **B)** Calprotectin levels, by using commercial or in-house ELISA. **C)** DNA isolated from equal volume of plasma samples from 16-week-old mice were tested for 16S rDNA levels by qPCR. **D)** Sixteen-week-old male and female mice were given FITC-dextran (5mg/mouse) by oral gavage, euthanized after 2h, and plasma samples were examined for FITC-Dextran concentration against control samples spiked with known concentration of FITC-Dextran. Value from the mouse with lowest plasma FITC-dextran level was considered as 1 for calculating relative values for rest of the mice within each cohort. **E)** cDNA preparations from the distal colon (left panel) and distal ileum (right panel) of 16-week-old mice were subjected to qPCR to assess the expression levels of key immune and tight-junction function proteins. Expression levels of individual factors were calculated against beta-actin values and used for generating the heat-map. **F)** RNA prepared from the distal colon of 16-week-old male and female mouse littermates were subjected to bulk-RNAseq and the gene counts of data were then normalized and subjected to visualization and statistical analysis using iDEP2.01 platform as described for Fig. 2. KEGG Pathway enrichment analysis of DEGs for significantly up- and down- regulated metabolic or signal transduction pathways in females compared to males. Additional analyses of this data set are shown in supplemental Fig. 3. *p-*values by Mann-Whitney test (comparison of males and females within specific age groups) for panels A-E. *<0.05, ***0.001.

*Gut microbiota impacts disease progression differently in NZM2328 males and females.* Previously we reported that the adult age, but not juvenile age, gut microbiota composition and the microbiota-mediated modulation of disease progression are different in males and females of lupus-prone SNF1 littermates [17]. Here, we examined fecal pellets from littermates of NZM2328 mice at 16 weeks of age for microbial community profiles. Principal component (PC) analysis found gender specific β-diversity clustering of gut microbiota with fecal samples of male and female NZM2328 mice **(Fig. 4A)**. On the other hand, **Fig. 4B** shows that α-diversity / species richness was not significantly different, statistically, in these male and female littermates. Compilation of the 16s rRNA gene sequences to different taxonomical levels showed that fecal microbiota in females had significantly higher abundance of Bacteroidetes phylum members **(Fig. 4C)**. Further, **Fig. 4D** and **supplemental Fig. 4A** showed that abundance of multiple major and minor microbial species and the overall predictive function of microbiota are considerably different in male and female littermates at 16-weeks of age. Microbial community profile of another cohort of male and female littermates tested at 8-weeks of age showed only modest differences in the fecal microbiota features **(supplemental Fig. 4B)**, relative to that of mice at 16-weeks of age **(Fig. 4A-C)**. These observations suggest gradual age and gender- specific changes in the gut microbiota of same maternal origin. Overall, these observations in NZM2328 mice were consistent with our previous observations from studies using the SNF1 mouse model, an F1 strain model of lupus with similar female bias [17].

**Fig. 4:**
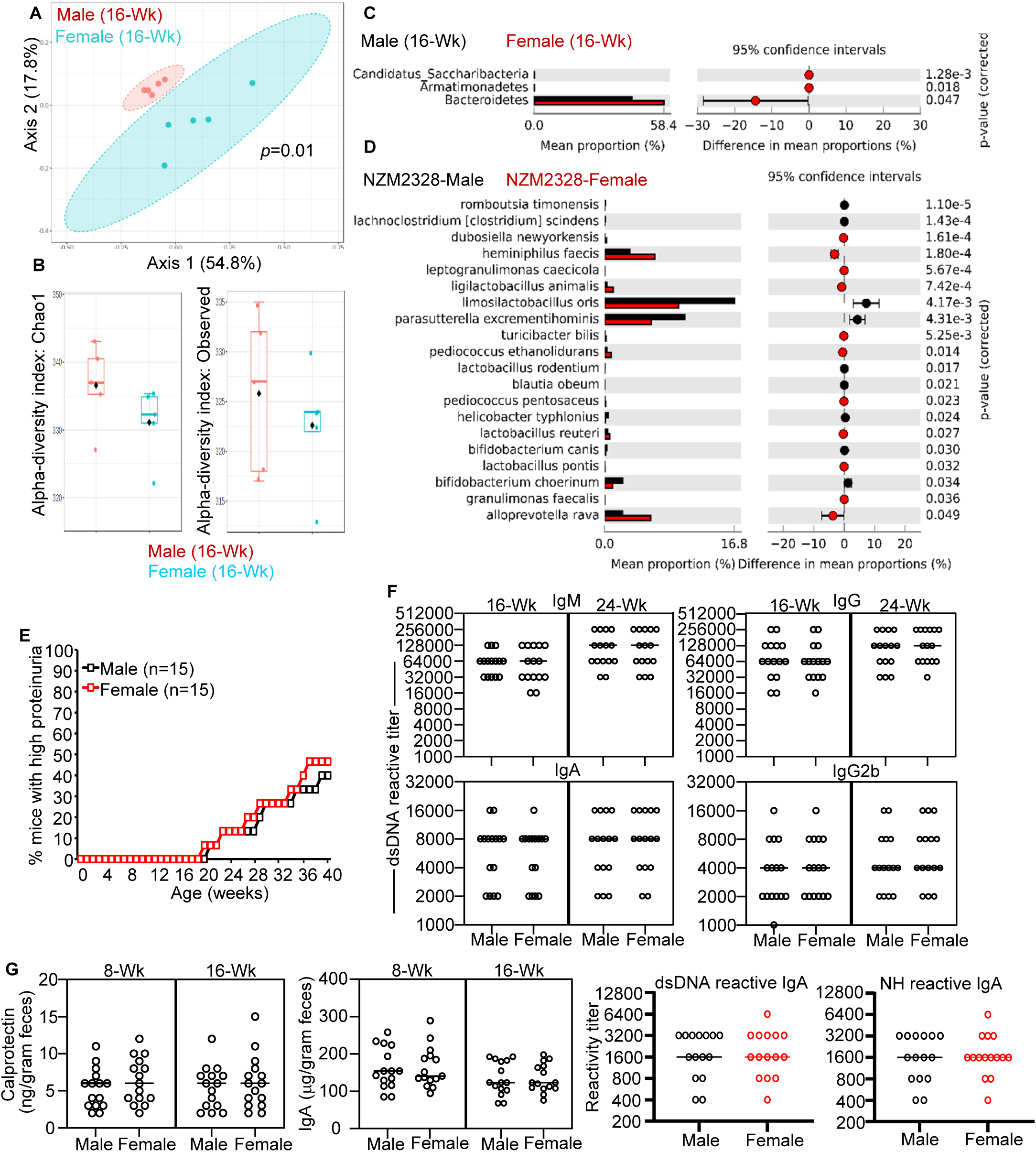
Gut microbiota composition and the impact of microbiota depletion in male and female NZM2328 mice. Fresh fecal pellets were collected at 16-weeks-of-age from male and female littermates (n=5) and DNA preparations of the fecal pellets were subjected to 16S rRNA gene (V3/V4 region) -targeted sequencing. The sequences were compiled to different taxonomical levels by employing 16S Metagenomics application of Illumina BaseSpace hub and the data was visualized by employing MicrobiomeAnalyst and STAMP applications. **A)** Principal component analysis (PCA) plots of samples representing β-diversity (Bray Curtis distance). **B)** α-diversity (Chao1 and observed) comparisons. **C)** Mean relative abundances of 16S rRNA gene sequences in fecal samples at phyla level. **D)** Mean relative abundances of sequences representing major and minor microbial communities at species level. Statistical analyses were done by employing two-sided Welch’s *t-test* and the *p*-values were FDR corrected using Benjamini and Hochberg approach. This 16S rRNA gene amplicon sequence data based on predictive functional profiles are shown in supplemental Fig. 4A. **E)** Cohorts of male and female NZM2328 mice of the SPF facility were maintained on drinking water containing broad-spectrum antibiotic cocktail (Abx), starting at 4-weeks-of-age and monitored for high proteinuria. **F)** Serum samples collected at different ages from Abx-treated mice were subjected to ELISA to determine the dsDNA-reactive IgM, IgG, IgA and IgG2b titers. **G)** Extracts of fecal samples from different age groups of Abx-treated male and female mice were examined for Calprotectin levels, total IgA concentrations, and dsDNA and NH reactive IgA titers by using commercial or in-house ELISA. Log-rank test (for panel E) and Mann-Whitney test (for panels F&G) were employed to compare males and females within specific age groups.

We then examined if microbiota depletion impacts disease progression in NZM2328 mice. Male and female mice were left untreated or given broad-spectrum antibiotic cocktail (Abx) in drinking water, starting 4-weeks of age to deplete the gut microbiota and monitored for systemic autoantibody levels and the timing of elevated proteinuria. As shown in **Fig. 4E**, compared to mice with an intact gut microbiota (Fig. 1A), the overall timing of onset of proteinuria was profoundly delayed in female NZM2328 mice that were given Abx. While about 80% of control females developed severe nephritis by 40 weeks of age, only <50% of the microbiota-depleted mice showed a similar degree of disease during this monitoring period. Importantly, as compared to untreated mice (Fig. 1A), microbiota depletion had only a modest impact on the timing of proteinuria onset in male mice. Forty-week severe nephritis incidence, indicated by high proteinuria, was <30% in untreated males compared to about 40% in Abx-treated mice suggesting a modest increase in disease activity in males, in contrast to the observed disease suppression in females. Further, as shown in **Fig. 4F**, the serum dsDNA-reactive IgA, IgG and IgM, and IgG2b (or other sub-class; not shown) autoantibody levels were not significantly different between antibiotic-treated male and female mice. Similarly, serum NH-reactive antibodies, spleen sizes, kidney pathology, and immune factor gene expression profiles were not considerably different between microbiota-depleted male and female NZM2328 mice (not shown). Importantly, **Fig. 4G** shows that microbiota depletion also eliminated the gender-specific differences in the fecal calprotectin abundance and nAg-reactive IgA levels, which were observed in untreated male and female mice (Fig. 3). Overall, these results suggest that gut microbiota influences lupus-like disease progression differently in NZM2328 males and females and the female bias in lupus-like disease in this strain of mice is microbiota-dependent.

### Gender specific differences in disease incidence and autoantibody levels are largely absent in GF NZM2328 mice

To validate the outcome of studies using microbiota-depleted NZM2328 mice, the strain was subjected to germ-free (GF) derivation. After GF derivation, multiple generations of NZM2328 mice were monitored for 40 weeks of age for the timing of clinical stage disease onset (high proteinuria) and survival (end stage disease). Systematic bi-weekly monitoring of mice housed in GF isolators for proteinuria showed that >75% of the females and >70% of the male NZM2328 mice develop severe nephritis/proteinuria by the age of 40 weeks **(Fig. 5A)**, compared to about >80% females and <30% males housed in our conventional facility (Fig. 1A). Additional cohorts of mice monitored for the frequency of end-stage disease feature including whole-body edema, poor body condition score or high proteinuria in the GF isolators were about 68% for females and >62% for male NZM2328 mice by 40 weeks of age and log-rank test showed no statistical difference in the timing of severe nephritis or end stage disease between male and female GF NZM2328 mice.

**Fig. 5:**
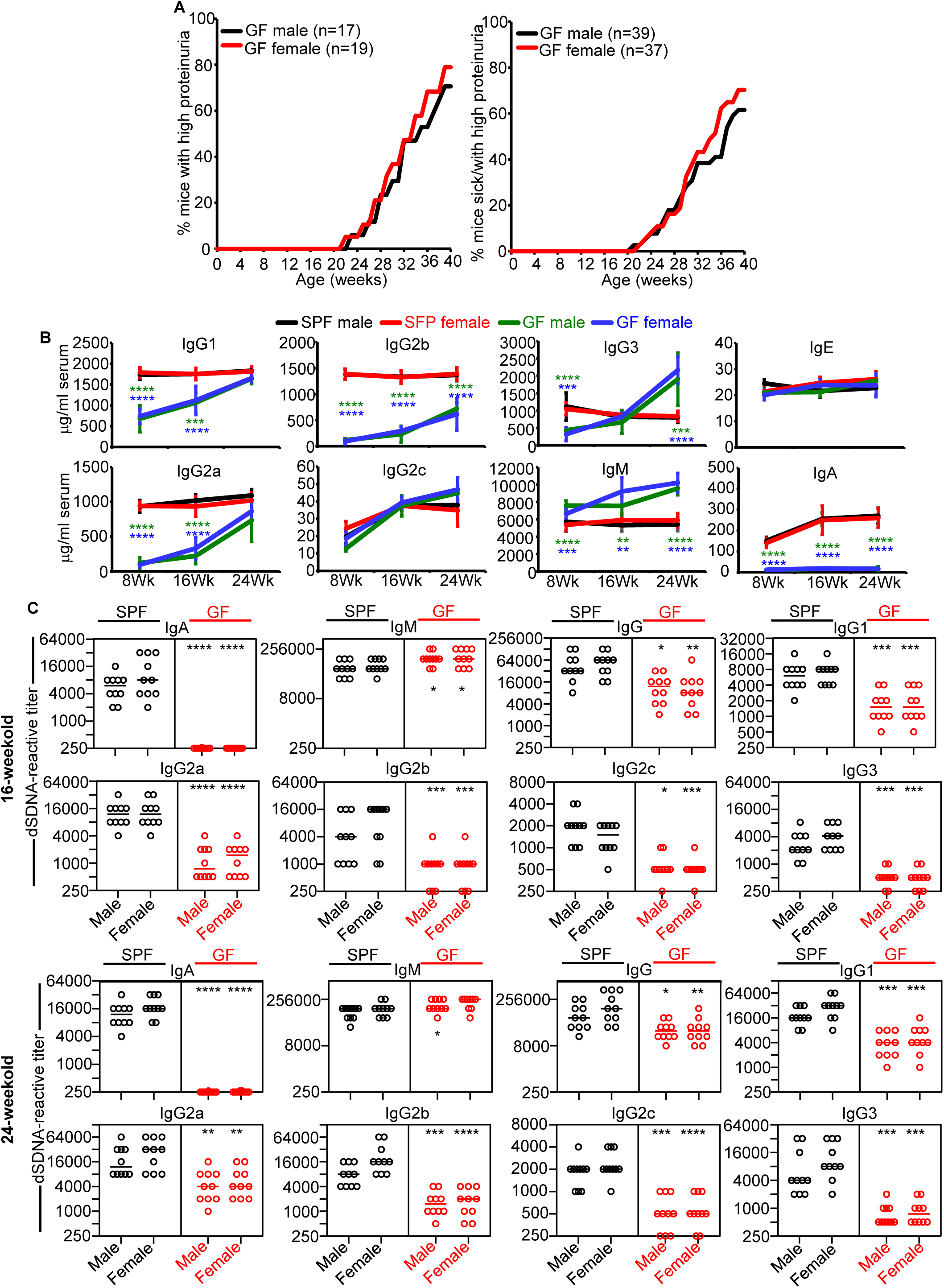
GF status eliminates gender difference in lupus-like disease incidence. **A)** Male and female GF NZM2328 mice were monitored for lupus-like disease features for up to 40 weeks of age. Percentage of mice with severe kidney disease based on high proteinuria values alone during biweekly testing (left panel) or on a combination of high proteinuria values and deteriorated body condition that required pulling mice from the gnotobiotic isolators (right panel) are shown. **B)** Serum samples collected from male and female GF and SPF mice (n=10/group) at different ages were subjected to antibody isotyping/subtyping by ELISA or multiplex assay and mean +/- SD of concentration values are shown. **C)** Serum samples collected from male and female GF and SPF mice (n=10 mice/group) at 16-weeks and 24 -weeks of age were subjected to ELISA to determine the dsDNA reactive titers of Ig isotypes and subclasses. Log-rank test (for panel A), Chi-squire test (for panel B), and Mann-Whitney test (for panels C&D) were employed to compare males and females within specific age groups. Specific age GF males and females were also compared to respective SPF counterparts for the *p-*values (*<0.05, **<0.01,***<0.001,****<0.0001) shown.

To compare the antibody production dynamics of NZM2328 mice in the absence and presence of microbiota exposure, abundances of serum Ig isotypes and IgG subclasses of GF and SPF NZM2328 mice at different ages were determined. **Fig. 5B** shows that both male and female GF mice produced significantly lower amounts of IgG1, IgG2a and IgG2b, especially at 8 and 16 weeks of age, as compared to their SPF counterparts. By 24 weeks of age, abundance of some of these IgG subclasses in GF mice were comparable to that of SPF mice. IgG3 subclass was produced at higher amounts by SPF mice at 8 weeks of age and GF mice at 24 weeks of age. While SPF mice showed similar levels of most IgG subclasses at different adult ages, GF mice showed a gradual age dependent increase in the abundance of these antibodies. GF mice showed persistently higher, and gradually increased, production of IgM isotype compared to SPF mice. Consistent with the previous reports on IgA production dynamics under GF conditions [40, 41], GF mice produced profoundly lower amounts of IgA isotype at all timepoints compared to their SPF counterparts. We have then compared the dynamics of circulating dsDNA and NH reactive IgA, IgM and IgG isotype (as well as IgG subtypes) in male and female SPF and GF NZM2328 mice starting at 8 weeks of age. Except for IgM, all isotypes (IgG and IgA) and IgG subclasses (IgG1, IgG2a, IgG2b, IgG2c and IgG3) of antibodies showed significantly lower nAg- reactivity titer in GF males and females at all time-points (16 and 24 weeks are shown) as compared to their SPF counterparts **(Fig. 5C&D)**. However, GF male and female showed comparable levels of serum nAg-reactive antibodies. Overall, nAg-reactivity titers of serum antibodies appear to be reflective of the relative abundance of most Ig isotypes and IgG subtypes shown in Fig. 5B.

### Gene expression profiles of SPF and GF NZM2328 mice reveal key microbiota-dependent differences

SPF male and female NZM2328 mice, but not their GF counterparts, showed significant differences in the timing of clinical stage disease onset and inflammatory features. Further, qPCR assay of cDNA prepared from the spleen tissues of male and female GF NZM2328 mice at different ages did not show a significant difference in the expression levels of key immune factors and tight junction proteins (not shown). Therefore, to determine microbiota-dependent, gender-specific early-age features pertinent to lupus-like disease, spleen and intestinal tissues from young adult (8-week-old) SPF and GF mice were compared for their gene expression profiles. Comparison of gene expression profiles of GF male and female did not show considerable differences (not shown). On the other hand, as shown in **supplemental Fig. 5A-C,** spleen tissues from 8-week-old SPF males and females showed a difference in the expression levels of specific genes and pathways, including NET and CCC, similar to the older adults of 16-week (Fig. 2), albeit at a relatively modest difference. Importantly, comparison of male SPF mouse spleen with male GF mouse spleen and female SPF mouse spleen with female GF mouse spleen revealed many common genes and pathways upregulated in both males and females in the presence of microbiota **(Fig. 6** and **supplemental Fig. 5D&E)**. While the key genes of some of these pathways such as *Mpo* and *Elane* are expressed at higher levels in both male and female SPF mice, difference in the expression levels of other neutrophil activation proteins such as S100a8, S100a9 and chaperone proteins Hspa1a, Hspa1b and Hsph1 are more pronounced in SPF females than in males compared to their GF counterparts **(Fig. 6A-B).** Among the key pathway genes upregulated in SPF male and female mice as compared to their GF counterparts, NET, CCC and IL17 signaling pathway genes are more pronounced in females than in males **(Fig. 6C-D)**. Genes of other key pathways such as platelet activation and ECM-receptor interaction were upregulated in both male and female SPF mice as compared to GF controls. However, only a limited number of genes and pathways were found to be expressed at higher levels in GF mice compared to their SPF counterparts (Fig. 6A-B).

**Fig. 6:**
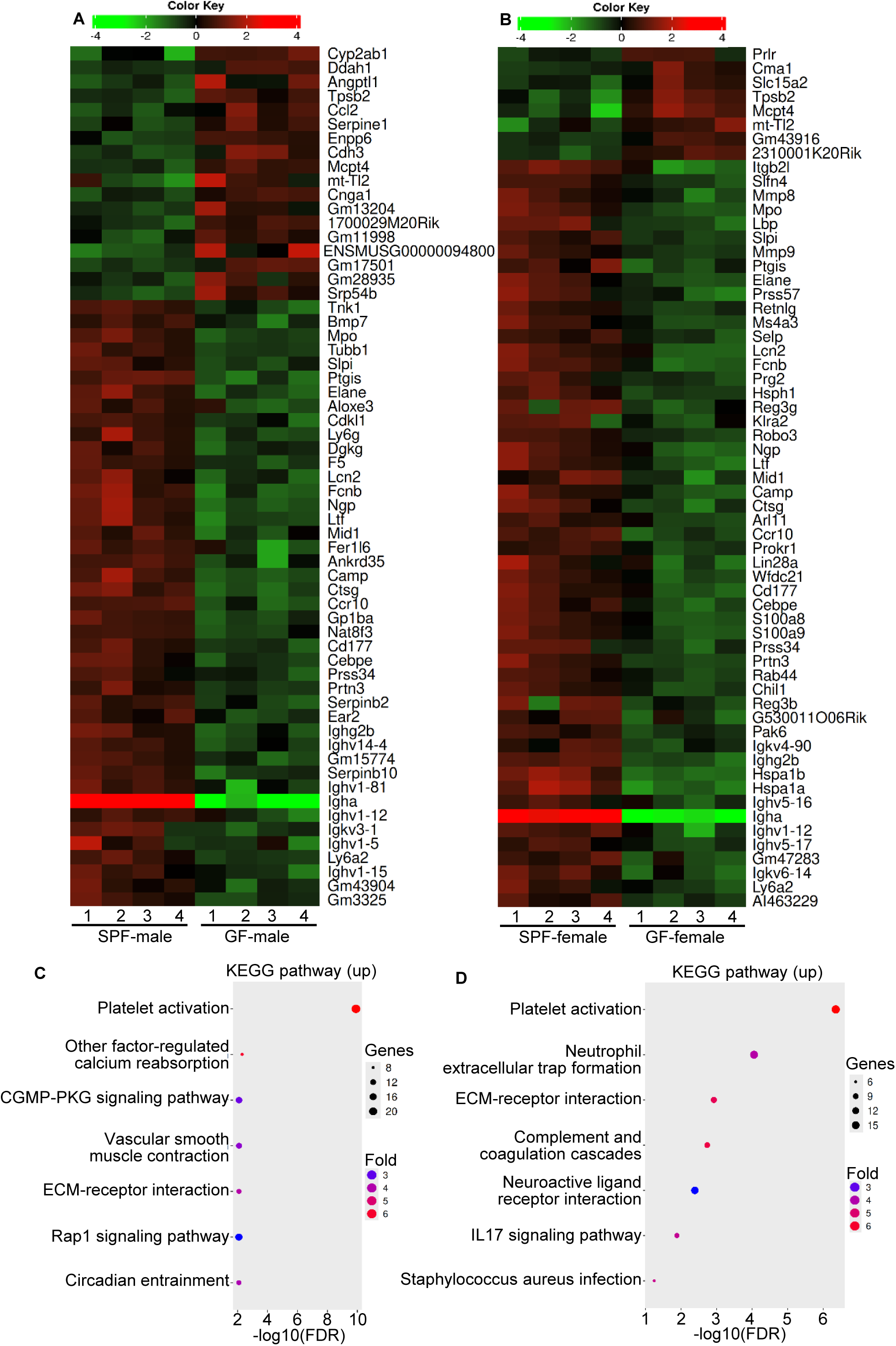
Microbiota exposure-associated activation of specific genes and pathways in the spleen are more pronounced in female. *NZM2328 mice.* RNA prepared from the spleen of 8-week-old male and female GF littermates and SPF littermates were subjected to bulk-RNAseq and the gene counts of data were then normalized and subjected to visualization and statistical analysis using iDEP2.01 platform as described for Fig. 2. Heatmaps representing major DEGs in individual samples: SPF male vs GF male **(A)** and SPF female vs GF female **(B)** are shown. KEGG Pathway enrichment analysis of DEGs for significantly up-regulated metabolic or signal transduction pathways in male SPF mice compared to male GF counterparts **(C)** and female SPF mice compared to female GF mice **(D)** are shown. Various analyses of 8-week-old male and female SPF mouse comparisons are shown in supplemental Fig. 6.

### Intestinal immune features were comparable in male and female NZM2328 mice under GF condition

Since male and female NZM2328 mice showed comparable systemic immune features, we examined if the intestinal immune features are any different in male and female GF NZM mice. qPCR assay of the distal ileum and colon tissues of male and female GF NZM2328 mice, unlike that of SPF mice (Fig. 3), did not show a considerable difference in the expression levels of key immune and tight-junction factors **(supplemental Fig. 6A)**. Fecal calprotectin levels and IgA features of SPF and GF males and females were compared to examine if there are gender-specific differences in these features in the absence of exposure to microbiota. As observed in **Fig. 7A&B**, calprotectin and IgA abundances and the overall nAg reactivity levels of IgA were profoundly lower in GF male and female mice as compared to their SPF counterparts. In fact, calprotectin was undetectable in the fecal samples of most GF male and female mice, even at 16 weeks of age. Similarly, in addition to the profoundly lower IgA levels, fecal extracts from GF NZM2328 mice showed no detectable dsDNA or NH reactive IgA irrespective of the age or gender. Comparison of gene expression profiles of male SPF mouse colon with male GF mouse colon and female SPF mouse colon with female GF mouse colon revealed large number common genes and pathways upregulated in both males and females when exposed to microbiota **(Fig. 7C and Supplemental Fig. 6C&D)**. Microbiota-dependent gender-specific differences in gene expression appears to be limited to a small number of pathways and genes such as Tnfrsf17, gzmb, ido1, Il22, Ccr4, Serpinb, etc. For examples, Tnfrsf17, gzmb, Serpinb and Ido1 are expressed at more pronounced levels in SPF female colon compared to GF female colon, Il22 and Ccr4 are expressed at higher levels in SPF male colon compared to their GF counterparts. Overall, these results along with the gene expression profiles of spleen tissues (Fig. 6) suggest that activation of lupus-like disease relevant inflammatory pathways in the systemic compartment of female mice is more pronounced than in their male counterparts in the presence of microbiota. However, comparison of the gene expression profiles of colon tissues from male and female GF mice showed no considerable differences (not shown). Overall, our studies using GF NZM2328 mice (Figs. 4-7) show that lack of exposure to microbiota eliminates gender bias in intestinal and systemic immune and non-immune features, pertinent to inflammation and autoimmunity. Further, although total and nAg-reactive IgA and IgG antibody levels are profoundly low in GF NZM2328 mice as compared to their SPF counterparts, 40-week disease incidence rate of GF males and females (Fig. 5A) were not significantly different from that of SPF females (Fig. 1A). On the other hand, SPF males showed lower 40-week disease incidence compared to their GF counterparts.

**Fig. 7:**
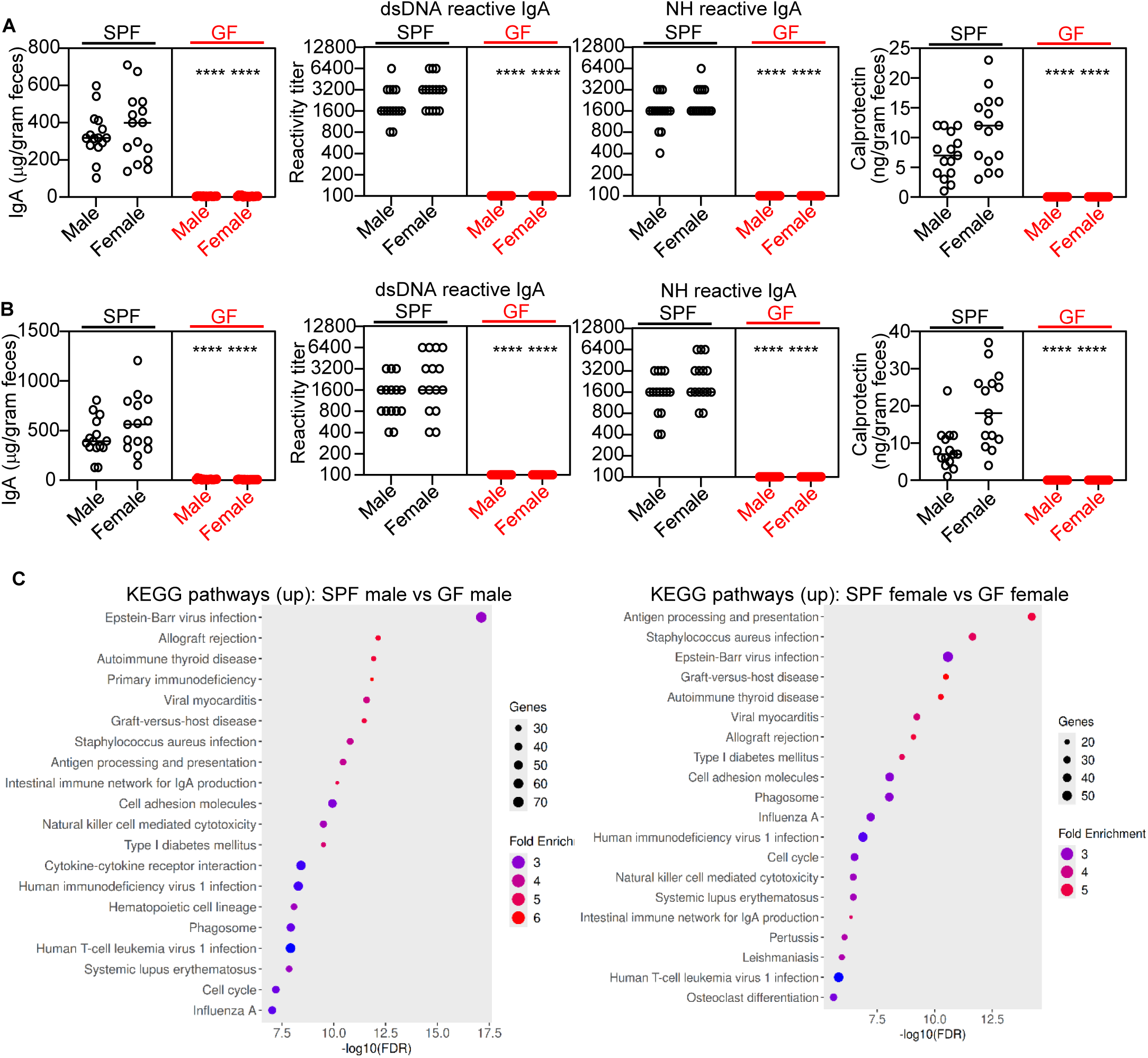
Lower neutrophil activity, and IgA and calprotectin production in GF-NZM2328 mice. A&B) Extracts of fecal samples from different age groups (8 and 16 week) of male and female SPF and GF NZM2328 mice were examined for Calprotectin and total IgA concentrations, and dsDNA and NH reactive IgA titers as described above. *p-*values by Mann-Whitney test for panels. **C)** RNA prepared from the distal colon of 8-week-old male and female GF littermates and SPF littermates were subjected to bulk-RNAseq and the gene counts of data were then normalized and subjected to visualization and statistical analysis using iDEP2.01. KEGG Pathway enrichment analysis of DEGs for significantly up-regulated metabolic or signal transduction pathways in SPF mice compared to their GF counter parts is shown.

### Conventionalization of GF NZM2328 mice restores gender differences in lupus-like disease associated features

Since GF NZM2328 mice showed little or no gender specific differences in autoreactive antibody levels and the timing of proteinuria onset and end stage disease, we examined if the gender bias can be restored in these mice upon microbiota conventionalization. Four-week-old male and female GF NZM2328 mice were pulled from GF isolators and conventionalized by housing on pooled, and equally distributed, dirty bedding from SPF NZM2328 breeding cages, microbial association in them was confirmed by anaerobic and aerobic culture fecal suspension as described for our previous report [32] (not shown). Fecal pellets collected from a cohort of conventionalized mice at 12-weeks of age (8-weeks post-conventionalization) showed differences in the composition of fecal microbiota **(Fig. 8A)** even though they were exposed to the same dirty bedding microbes. This suggests that gender-specific diversion in gut microbiota can occur in a short period, at young adult ages. To assess the impact of microbiota conventionalization of GF-NZM2328 mice on lupus-like disease progression, multiple batches of conventionalized mice were monitored for the timing of kidney disease indicated by high proteinuria. As observed in **Fig. 8B**, conventionalized NZM2328 mice, both male and females, showed delayed onset of proteinuria compared to their SPF and GF counterparts (Fig. 1A and Fig. 5A). The delay in the onset of severe proteinuria was more pronounced in males than in females. In addition to this delay in disease onset, a reduction in the 40-week disease incidence was observed in conventionalized male NZM2328 mice compared to their GF counterparts (Fig. 5A), but comparable to that of SPF mice (Fig. 1A). On the other hand, the rate of 40-week disease incidence of conventionalized female NZM2328 mice was similar to that of female SPF NZ2328 mice (shown in Fig. 1A). To determine the impact of gender influenced microbiota on lupus-like disease onset in NZM2328 mice, GF males and females were given fecal microbiota of the same or opposite gender SPF mouse donors of >16-weeks of age by oral gavage. **Fig. 8C** shows that fecal microbes of opposite gender have only a modest modulatory effect on the recipients, perhaps due to the gradual alteration of microbiota by host and environmental factors, post-conventionalization. Nevertheless, the disease-inhibitory effect of male donor fecal microbiota appears to be more pronounced than the disease-promoting activity of female donor microbes.

**Fig. 8:**
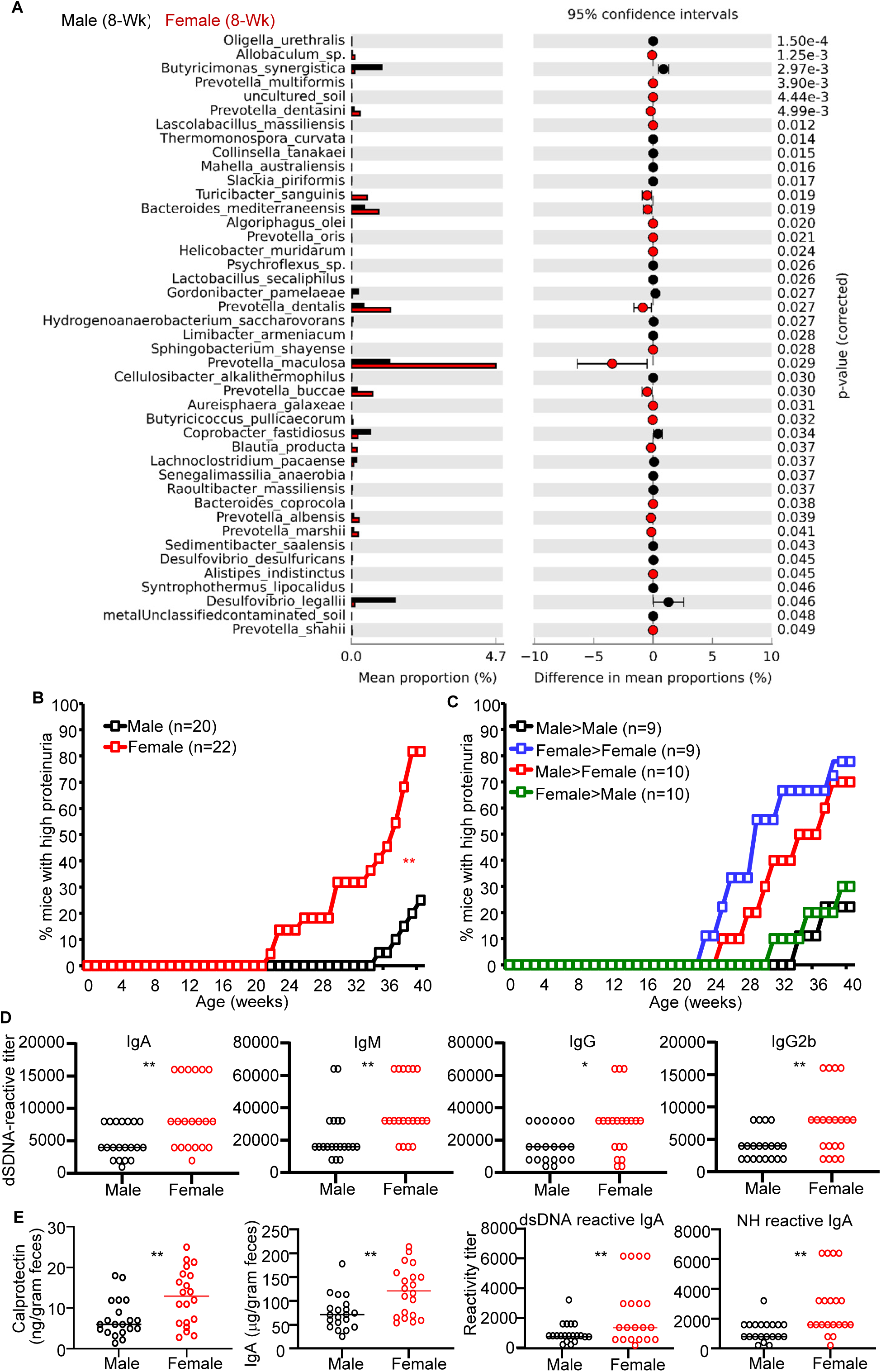
Microbiota conventionalization restored gender specific differences in microbiota and in lupus-like disease features. For microbiota conventionalization, four-week-old male and female GF NZM2328 mice were pulled from gnotobiotic isolators and housed on, pooled and equally distributed, dirty bedding from SPF NZM2328 mouse breeding cages. Bedding material was replaced with fresh dirty bedding, thrice, within a week. **A)** Fresh fecal pellets were collected from conventionalized representative male and female mice (n=4 mice/group) at 12 weeks of age (8-weeks post conventionalization) and subjected to 16S rRNA gene amplicon sequencing as described for Fig. 3. Mean relative abundances of sequences representing major and minor microbial communities at species level are shown. **B)** Conventionalized (ex-GF) male and female NZM2328 mice were tested for urinary protein values every week for up to 40 weeks of age. Percentages of mice with severe nephritis as indicated by high proteinuria (≥10mg/ml) are shown. **C)** Cohorts of GF male and female mice were conventionalized as described above, using dirty bedding from 16-week-old male and female mice as indicated. These mice, which also received fresh fecal suspension from the respective donor mice by a single oral gavage before housing on dirty bedding, were tested for proteinuria every week for up to 40 weeks. **D)** Serum samples collected at 16-weeks of age from microbiota conventionalized mice of panel B were subjected to ELISA to determine the dsDNA reactive IgM, IgG, IgA and IgG2b titers. **E)** Extracts of fecal samples from microbiota conventionalized mice of panel B were examined for Calprotectin levels, total IgA concentrations, and dsDNA and NH reactive IgA titers. Statistical comparison by log-rank test for panel B&C and Mann-Whitney test for panels D&E. *<0.05, **<0.01.

Microbiota-conventionalized mice were also tested for serum nAg-reactive antibodies, and the abundance and nAg-reactivity of fecal IgA as well as levels of inflammation marker calprotectin at 16-weeks of age. As expected, conventionalization caused an increase in the dsDNA and NH reactive serum IgA, IgM, IgG and IgG2b levels in both male and female mice **(Fig. 8D)**, as compared to their GF counterparts (Fig. 5). Consistent with the serum antibody levels, fecal IgA abundance and nAg reactivity were significantly higher in conventionalized females as compared to males **(Fig. 8E)**. Further, fecal calprotectin levels were also found to be higher in conventionalized female NZM2328 mice as compared to their male counterparts. Importantly, serum and fecal dsDNA and NH reactive antibody titers were not only higher in conventionalized female mice, but also the gender specific differences in the autoantibody levels were more pronounced in conventionalized mice than in what was observed in SPF mice (Fig. 1C&3A). Of note, comparison of serum dsDNA and NH reactive antibodies as well as fecal IgA features between cis-gender and trans-gender fecal microbiota transfer recipients of Fig. 8C showed no statistically significant differences (not shown). Nevertheless, collectively, these results confirm that the gender bias in lupus-like disease in NZM2328 mice is microbiota-dependent, and it is primarily a disease protective effect in males rather than the influence on females contributing to the disease-associated sex dimorphism. Previous studies including ours using other autoimmune models [17, 25, 42] have shown that the microbiota increases testosterone levels and testosterone-influenced microbiota suppresses disease progression in males. We found that testosterone levels in adult GF NZM2328 mice were modestly lower and restored to a level similar to that of their SPF counterparts upon conventionalization (not shown) indicating a potential contribution of testosterone-microbiota interaction to microbiota-dependent protection of males from the lupus-like disease.

### Conventionalization of GF NZM2328 mice restores the female bias in specific gene expression pathways

Studies described above show that SPF male and female NZM2328 mice, but not their GF counterparts, presents gender specific differences in the gene expression profiles related to pro-inflammatory pathways (Fig. 6 and supplemental Fig. 5). Therefore, to assess if microbiota introduction influences genetic pathways differently in male and female NZM2328 mice, spleen gene expression profiles of conventionalized mice were determined. **Fig. 9A** shows that a large number of genes are differentially expressed in male and female NZM2328 mice conventionalized with/exposed to the same source fecal microbes within 8 weeks of conventionalization. The numbers of genes showed increase in expression upon conventionalization were higher in females as compared to males. Many of the genes and pathways overrepresented in SPF females as compared to SPF males (Fig. 2) and young SPF females as compared to age-matched GF females (Fig. 6) were also upregulated in conventionalized females as compared to male counterparts **(Fig. 9B)**. For instance, Kegg pathway and GO analyses of DEG show NET, platelet activation, ECM-receptor interaction, coagulation, and IL17 signaling pathway genes are expressed at higher levels in conventionalized females compared to their male counterparts. Moreover, neutrophil-associated calprotectin protein genes (S100a8 and S100a9) were expressed at higher levels in conventionalized females compared to their male counterparts. Overall, these results show that microbial factors contribute to gender-specific differential activation of inflammatory and anti-inflammatory pathways that modulate lupus-like disease progression in NZM2328 mice.

**Fig. 9:**
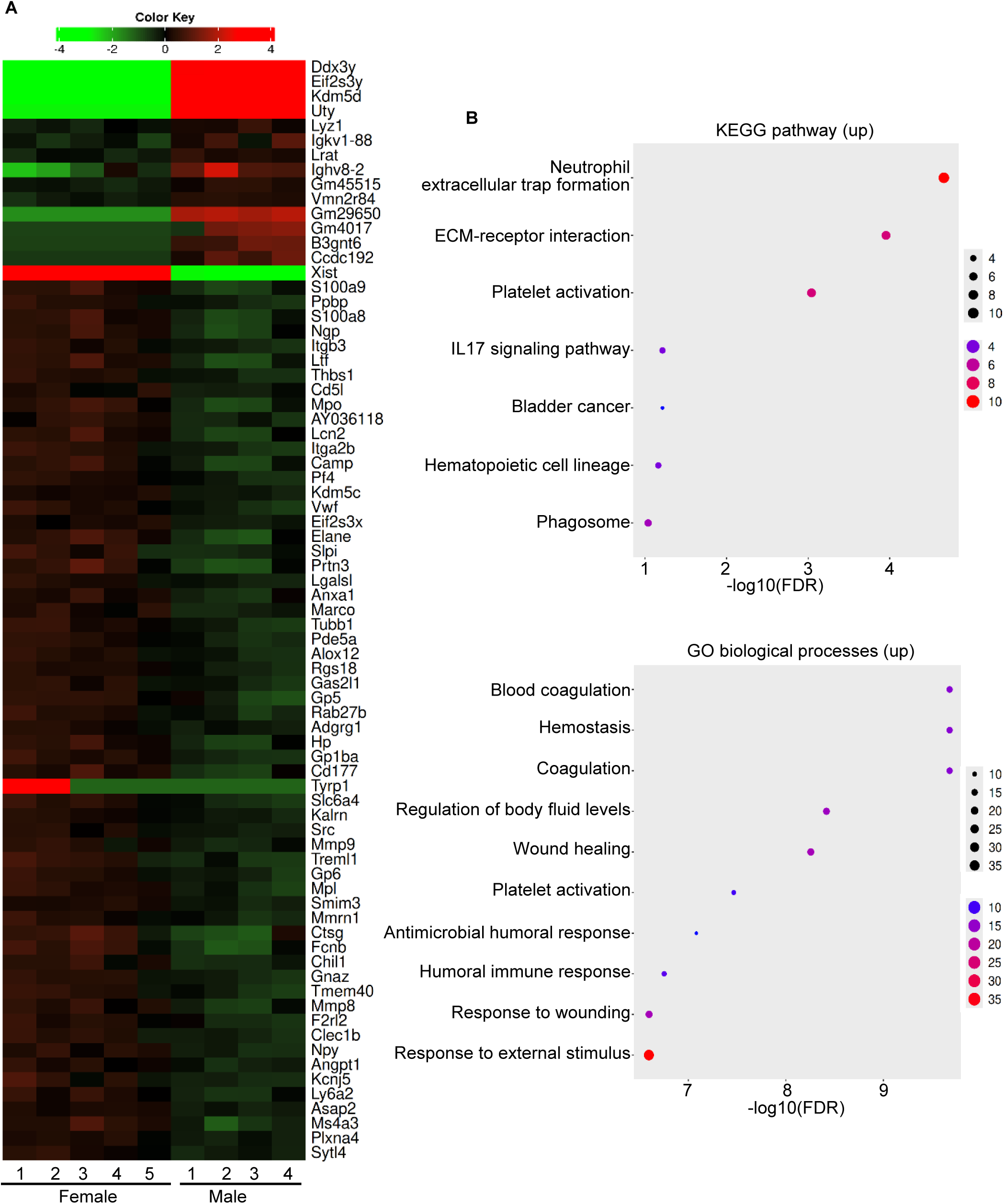
Microbiota conventionalization restored gender specific differences in the spleen gene expression profiles. RNA prepared from the spleen of 12-week-old microbiota-conventionalized male and female NZM2328 mice (as described for Fig. 8A&B) were subjected to bulk-RNAseq and the gene counts of data were then normalized and subjected to visualization and statistical analysis. **A)** Heatmap representing major DEGs in individual samples. **B)** KEGG Pathway enrichment and GO analyses of DEGs for significantly up-regulated pathways in females compared to males.

## Discussion

Gender bias in disease incidence is prevalent in some autoimmune diseases, particularly in SLE with up to 9:1 female to male ratio [43–45]. The influence of sex hormones on lupus autoimmunity has been studied [46–49]. Previous reports by others that used non-obese diabetic mouse model of T1D have linked testosterone-influenced gut microbes to gender bias in autoimmunity [25, 26]. Our previous report [17] has shown that gut microbiota contributes differently to lupus-like disease in SNF1 mice with male mice showing a protective effect and orchidectomy partially eliminating this disease-protective effect of microbiota. Our previous studies [17, 30] also showed that the gut mucosa of lupus-prone female SNF1 mice have more inflamed phenotype and depletion of gut microbes suppresses the intestinal pro-inflammatory features and systemic autoimmune progression in them. Further, it has been shown that lupus-and T1D- prone models in GF background do develop disease, indicating that exposure to microbes is not required for autoimmune progression, and the host genetics determines the development of these diseases and gut microbiota, in association with host factors, primarily modulates the rate of disease progression [22–26, 42]. However, systematic studies to examine the influence of gut microbiota on female bias in lupus-like disease and systemic autoimmune progression by employing human SLE relevant gnotobiotic animals are still lacking. Here, studies using NZM2328 mouse model, a slow progressing disease model which presents profound gender bias in the onset of clinical stage disease and employing microbiota depletion and GF derivation as well as microbial association strategies, we show that microbiota influences lupus-like disease outcomes differently in males and females. Consistent with our previous observations [17] using microbiota depleted and castrated SNF1 mice, here we found that microbiota has a minor disease-promoting effect in females, but it suppresses the disease progression in males significantly. Importantly, our current study also shows that microbiota-and host factor-dependent: 1) overactivation of specific immune pathways such as NET and CCC pathways in females and 2) suppressed pro-inflammatory pathways in males could be responsible for the female bias of lupus-like disease in NZM2328 mice.

Female NZM2328 mice housed in the SPF facility show profound overall higher disease susceptibility, as indicated by the timing of proteinuria onset and 40-week disease incidence, compared to their male counterparts. However, as observed before [19, 28], the circulating nAg-reactive antibody levels were not significantly different in males and females until very close to kidney disease/severe proteinuria onset age. This suggests that factors other than the autoantibody abundance are also contributing to the gender specific differences in disease progression in these mice. In addition to many known pro-inflammatory cytokines and X-chromosome linked genes including TLR7, TLR8 and CXCR3 that can contribute to lupus pathology, hyperactivation of NET, CCC, platelet and ECM-receptor interaction pathways appear to dominate in SPF females at pre-nephritis age as compared to their male counterparts. While these pathways have been implicated in various autoimmune diseases, particularly SLE and arthritis [50–57], their involvement in lupus-associated female bias and the microbiota dependency are not fully understood. In this regard, neutrophils isolated from female SLE patients tend to show higher NETosis properties as compared to these cells from men [54].

NETosis, CCC, platelet and ECM-receptor interaction pathways are not independent functional entities but intensive interactions between these processes have been reported [58, 59]. NETosis process leads to tissue damage by driving inflammation, promoting autoantibody formation, and causing immune complex deposition [60, 61]. It has also been shown that, within this inflammatory milieu, ROS primes and NETs activate inflammasome in immune cells causing release of cytokines such as IL-1β and IL-18 [62, 63]. These cytokines can further amplify ROS accumulation and NETosis, establishing a feed-forward inflammatory loop and progressive tissue damage in clinical conditions like SLE.

Our previous studies using SNF1 mice and human stool samples have shown that inflammatory features of gut mucosa is a key feature of SLE, particularly the female bias [17, 19, 21, 30]. Fecal proteomics, cytokine expression profiling, calprotectin and IgA analysis studies revealed that gut mucosa of female SNF1 mice, which presents female bias in systemic and mucosal autoantibody levels and disease characteristics, have elevated inflammation as early as at juvenile as compared to their male counterparts[17, 30]. Here, we found that while fecal IgA features are comparable in male and female NZM2328 mice at pre-clinical disease ages, neutrophil-associated gut inflammation marker calprotectin was produced at higher levels even in younger adult females as compared to their male counterparts. Pro-inflammatory gene expression profiles of intestinal tissues, colon particularly, reiterates the notion that immune-mediated events of gut mucosa have a role in female bias of lupus.

Our observation that the gut microbiota composition of male and female littermates of NZM2328 strain differs at adult age is in agreement with previous reports including ours that used different autoimmune models [17, 25, 26, 42, 64, 65]. These reports suggested that the same maternal microbiota acquired at young ages by male and female litters could be influenced by host factors such as hormones and environmental factors and can change during adult ages and influence the disease outcomes [5, 17, 25, 42, 66–71]. Similar to what was observed in SNF1 mouse model [17], microbiota depletion using Abx starting at 4 weeks of age resulted in suppression of systemic autoantibody levels, fecal calprotectin and IgA abundance, nAg reactivity of IgA and 40-week disease incidence in female, but not in male, NZM2328 mice. In fact, a modest increase in disease activity was observed in microbiota-depleted males indicating opposite effects of gut microbiota on lupus autoimmunity in females and males. This, along with our previous report employing microbiota-depleted SNF1 mice [17], suggests that the impact of gender-microbiota interaction on disease outcomes is consistent across different lupus-prone mouse models with substantial female bias.

Gnotobiotic studies using T1D model have shown that gender bias in autoimmune disease onset profoundly diminished in the absence of exposure to microbiota and the microbiota-sex hormone, particularly testosterone, interaction contributes to gender bias to some degree [25, 42]. Employing microbiota-depletion and orchidectomy approaches, we found similar microbiota-testosterone interaction-associated protection of male SNF1 mice, as compared to their female counterparts, from lupus-like disease [17]. Here, GF NZM2328 mice show similar elimination of female bias in lupus-like disease in the absence of gut microbiota, with male and female mice having comparable serum autoantibody levels and extremely low fecal IgA and calprotectin levels at different ages, and the 40-week disease incidence. Importantly, although host genetics, but not microbiota, appear to be the primary factor responsible for initiating autoimmunity in these mice, GF mice showed significantly lower circulating autoreactive antibody levels as well as fecal IgA abundance and nAg reactivity, and fecal calprotectin levels compared to their SPF counterparts. This suggests that gut microbiota contributes to nAg-reactive antibody production in the systemic and mucosa compartments, irrespective of their pathogenicity, as well as the inflamed gut mucosa phenotype. This suggests that gender difference in the timing of disease onset in this model is potentially contributed by microbiota-host factor interaction dependent accelerated or suppressed pathogenic autoantibody and other pro-inflammatory factor production in females and males respectively.

As indicated by the lack of significant differences in the nAg-reactive antibody levels, autoantibody levels alone do not appear to be the primary factor in disease process in lupus-prone NZM2328 mice. Our observations that: 1) biological pathways such as NET formation, CCC and platelet, and ECM-receptor interaction are mostly activated only when exposed to microbiota, 2) NET formation and CCC gene activation is more pronounced in females than in males in the presence of gut microbiota, and 3) these pathways are activated in both male and female SPF mice, but other confounding host factors impacted by microbes may be suppressing the disease process in males are all highly relevant in this regard. Our observations suggest that the inherent susceptibility of females to microbiota-dependent hyperactivation of these pathways compared to males may be, in part, determining the female bias in lupus-like disease outcomes. Our observations that the female bias in activation of these pathways as well as disease-associated differences are lost up on GF derivation and restored in GF mice upon microbiota conventionalization validate this notion.

Multiple observations: 1) SPF female and GF male and female NZM2328 mice showed somewhat comparable disease progression, 2) systemic autoantibody dynamics is not profoundly different in SPF males and females particularly at younger ages, 3) both male and female GF NZM mice produce profoundly lower amounts of most antibody isotypes and IgG subtypes, and the overall autoantibody titers are lower in them, and 4) fecal levels of neutrophil-associated inflammation marker calprotectin are elevated in females, microbiota dependently also point to the involvement of complex interaction between multiple factors such as antibody abundance and function, complement and neutrophil activation, and various other host factors including sex hormones that are differentially expressed in males and females in the disease process. It is possible that differences in the composition of gut microbiota which is influenced by the gender associated host-factors such as sex hormones may be impacting the activation of NET and CCC pathways differently in male and female NZM2328 mice. It is also possible, as reported in the context of various immune cell populations [72] [12–16, 73, 74], that testosterone and estrogen may have differential effects on neutrophils leading to female bias in disease outcomes. Hence, systematic studies are needed in the future to determine the impact of sex hormones on microbiota-dependent activation of neutrophils and CCC pathways.

In conclusion, our study demonstrates that gut microbiota-dependent differential activation of various immune regulatory and pro-inflammatory pathways in male and female mice contributes to female bias in lupus-like disease outcome in a human SLE relevant mouse model. While host genetic factors are the primary cause of autoimmunity in this model, gut microbiota and host factors shaped microbiota composition could modulate the disease outcomes significantly where the primary effect is disease protection in males rather than disease acceleration in females. Therefore, additional comprehensive mechanistic studies are needed in the future to determine the contribution of gender specific, sex hormone and microbiota dependent, NET and CCC and other related pathway activation in female bias in SLE. Importantly, it is also important to distinguish the pathogenic and disease-protective components of pathways including NET formation, CCC and platelet, and ECM-receptor interaction and study how host-factors such as sex hormones are impacting the activation of such pathways by gut microbiota.

## Footnote

This work was supported by unrestricted research funds from MUSC and National Institutes of Health (NIH) grant R21AI136339, 5R01AI138511 and R01DK136094. This project was supported in part by the Gnotobiotic Animal Core, which receives funding from the MUSC COBRE in Digestive and Liver Disease, NIH P20GM130457. Dr. Vasu is the guarantor of this work and, as such, has full access to all the data in the study and takes responsibility for the integrity of the data and accuracy of the data analysis. The authors are thankful to Cell and Molecular Imaging, Pathology, immune monitoring and discovery, and flow cytometry cores of MUSC for the histology service, microscopy, FACS and multiplex assay instrumentation support.

