## Supplemental Figures for "Neutrophil Extracellular Trap Formation and Complement Activation Pathways Dominate Microbiota-dependent disease bias in lupus-prone female NZM2328 mice"

**Fig. S1**

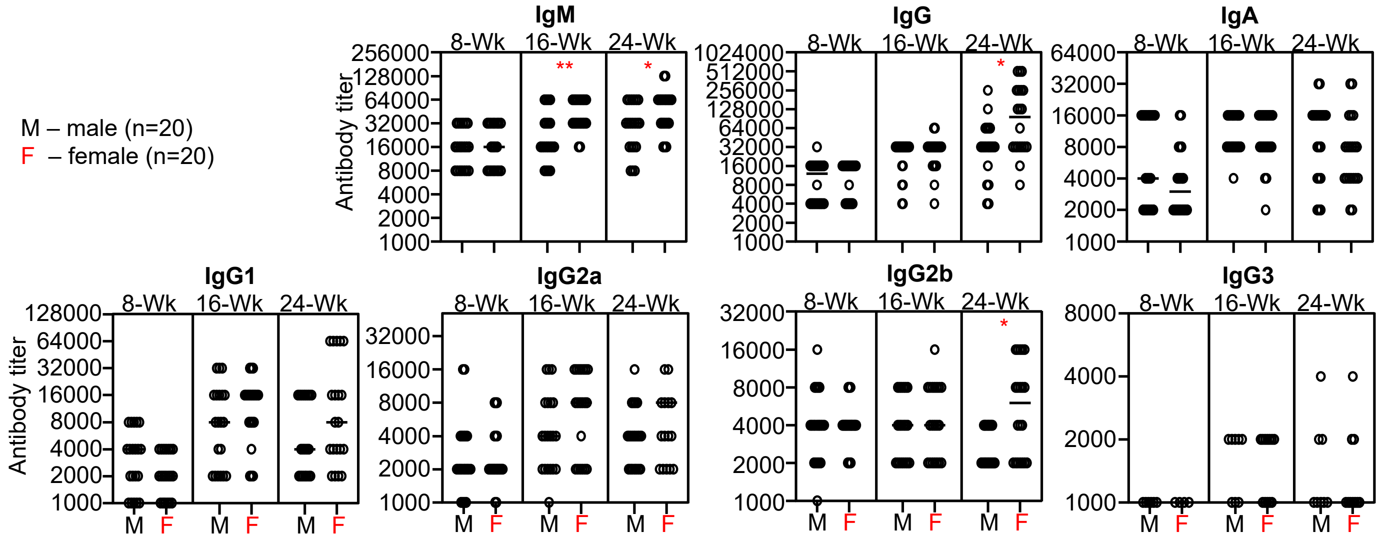


*Gender specific differences in NH reactive antibody levels in male and female NZM2328 mice housed in an SPF facility.* Serum samples collected from male and female mice at different ages (described in Fig. 1) were subjected to ELISA to determine the NH reactive titers of Ig isotypes and subclasses. p-values by Mann-Whitney test (comparison of males and females within specific age groups) *<0.05, **0.01.

**Fig. S2**

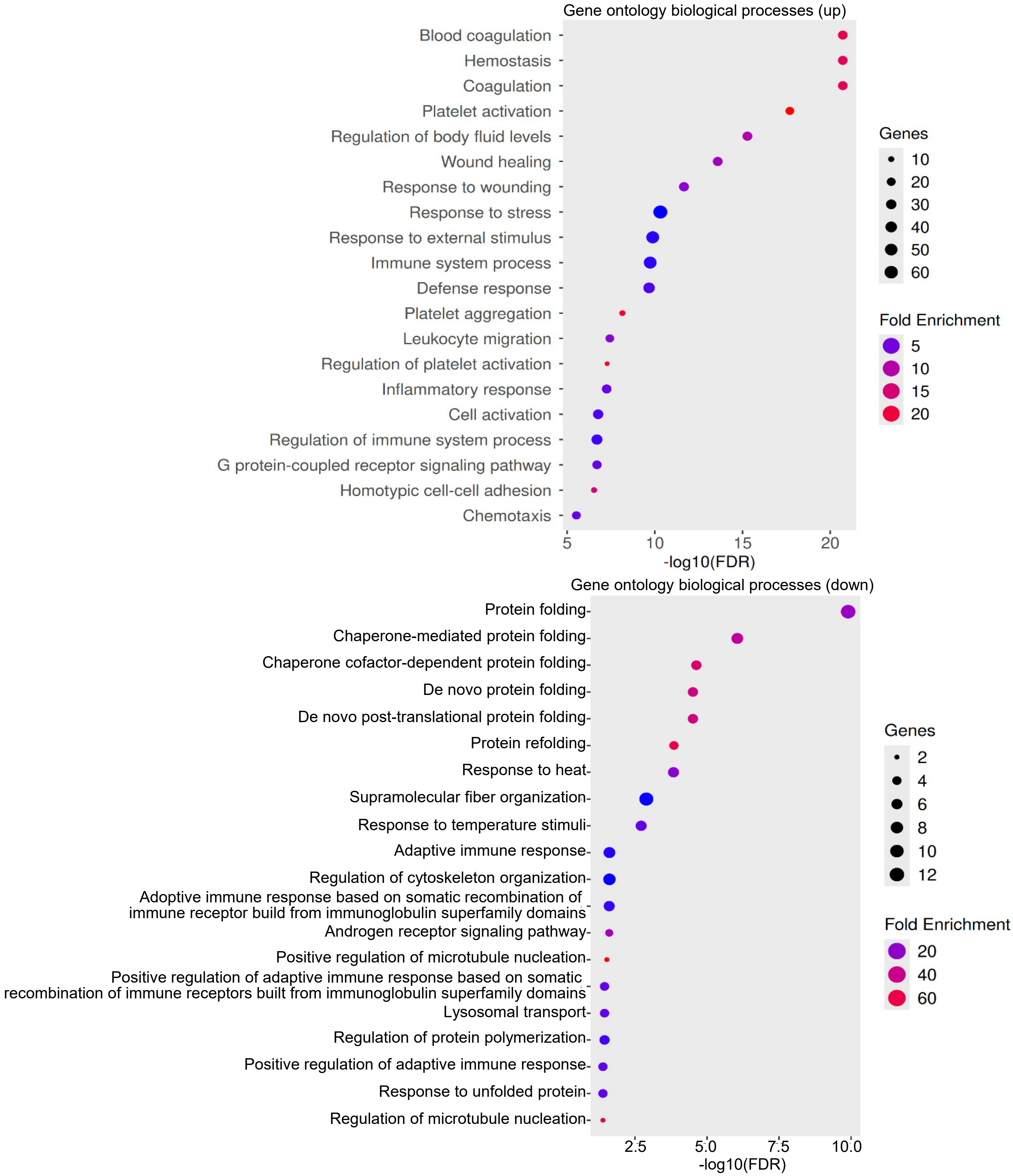


*Differences in gene expression profiles of the spleen tissues from male and female NZM2328 littermates.* Bulk RNAseq of spleen tissues of 16-week-old male and female NZM2328 mice were done as described for Fig. 2 GO enrichment analysis of significantly up- and down- regulated biological processes in females as compared to males is shown.

**Fig. S3**

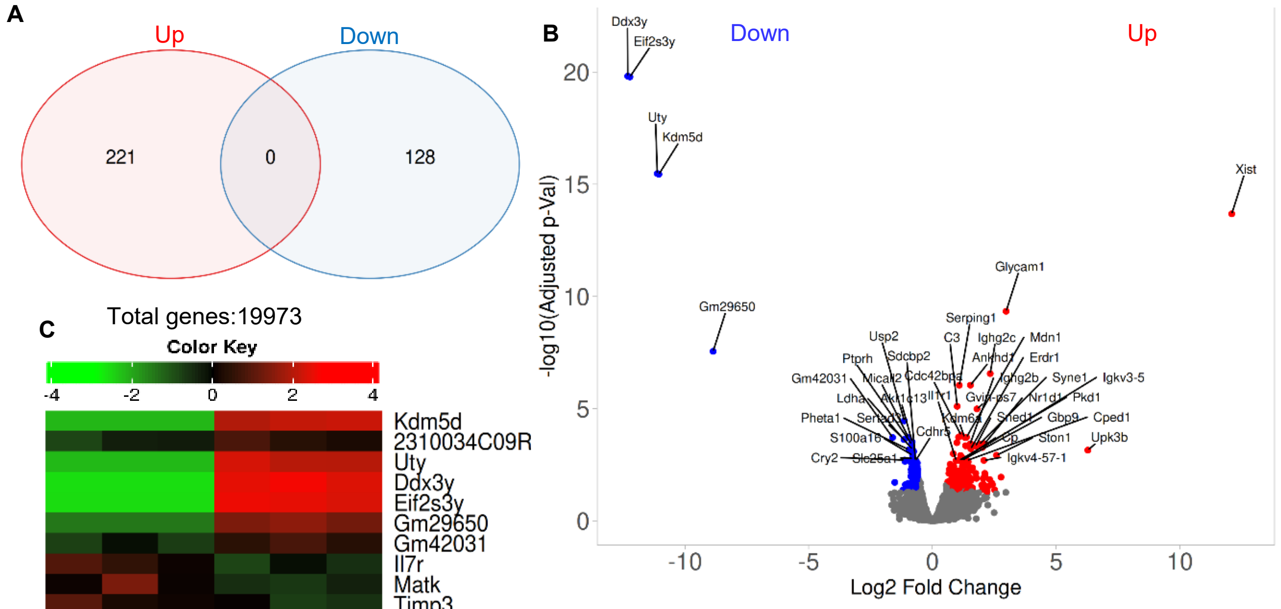


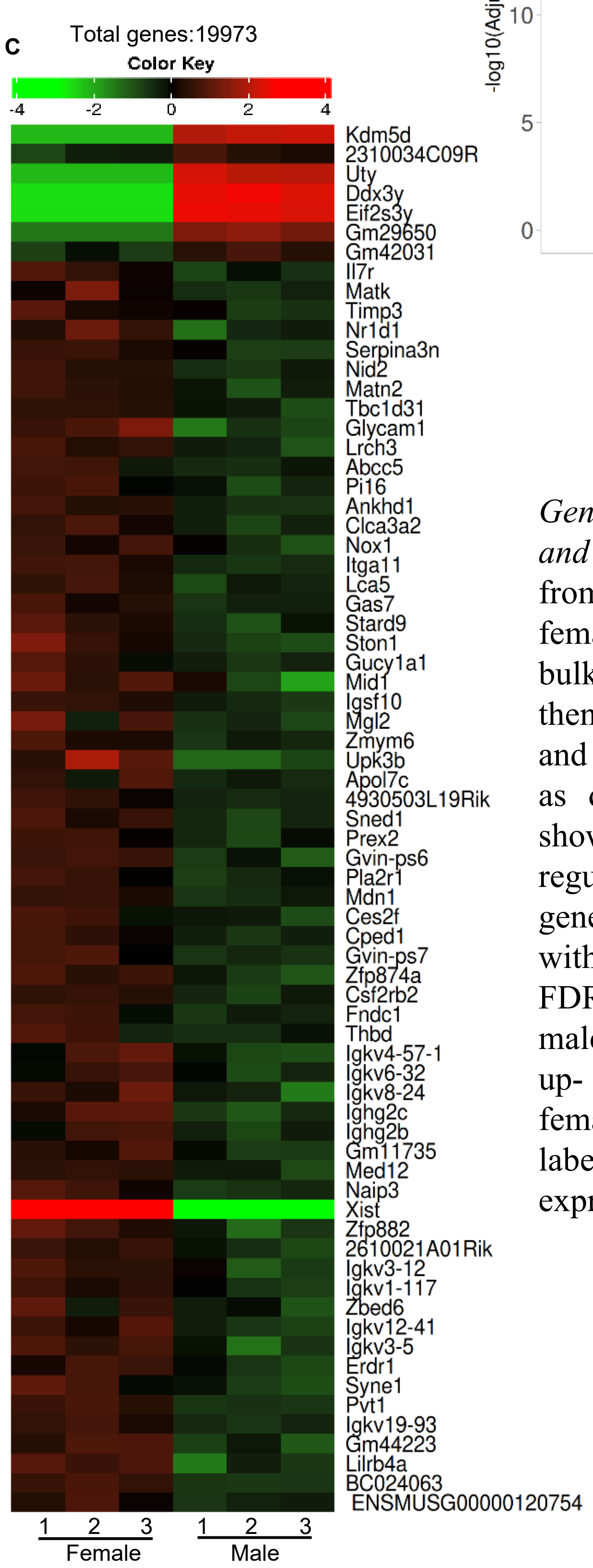


*Gene expression profiles of 16-week-old male and female SPF NZM2328 mice.* RNA prepared from the distal colon of 16-week-old male and female mouse littermates were subjected to bulk-RNAseq and the gene counts of data were then normalized and subjected to visualization and statistical analysis using iDEP2.01 platform as described for Fig. 3. **A)** Venn diagrams showing number of significantly up- and down-regulated genes among differentially expressed genes (DEGs; identified employing DESeq2 with a minimum fold change of 1.5-fold and FDR cutoff as 0.05) in females compared to males. **B)** Volcano plot showing significantly up- and down- regulated genes among DEGs in females compared to males with top genes labeled. **C)** Heatmap showing top differentially expressed genes in females and males.

Fig. S4

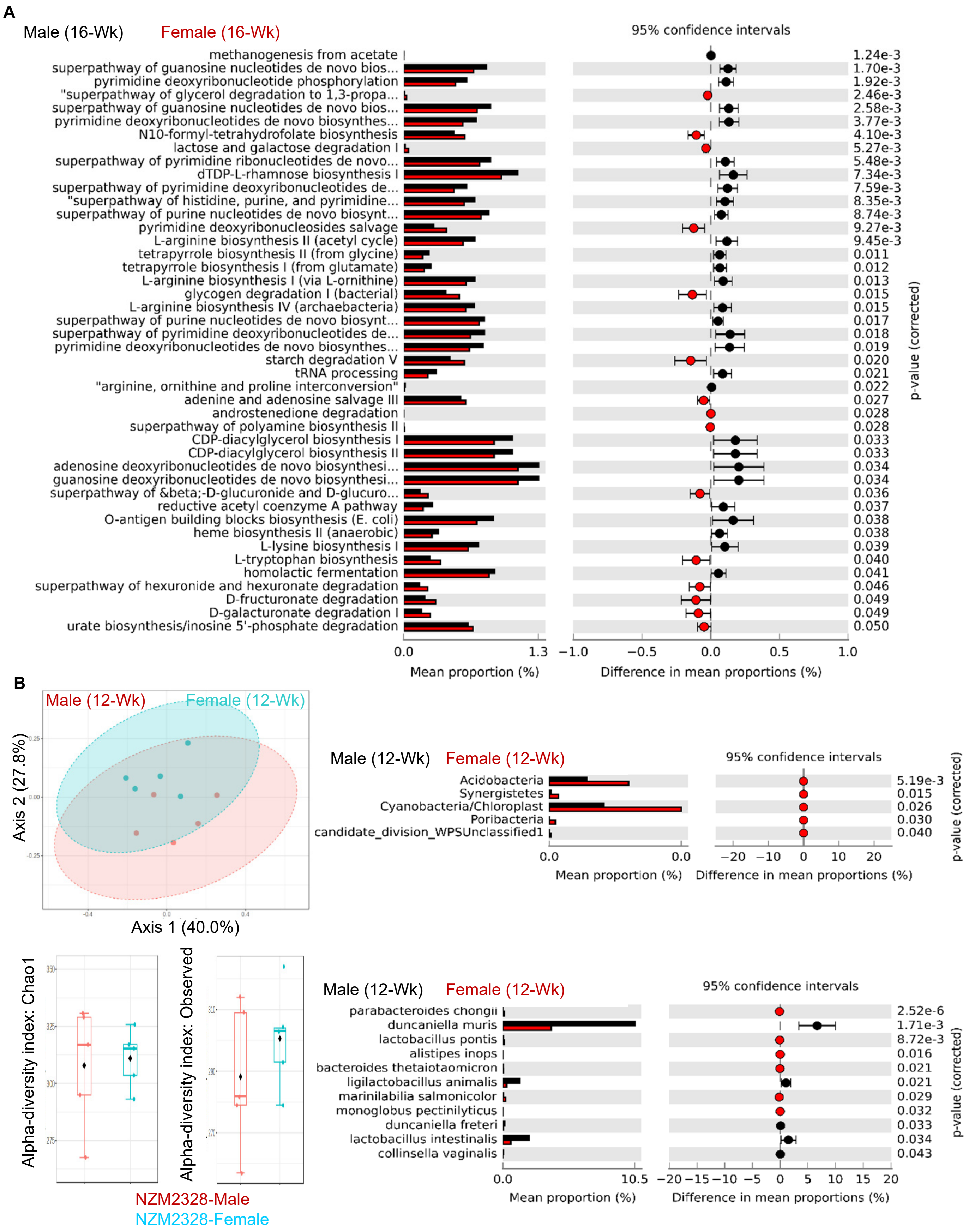
A
Male (16-Wk)
Female (16-Wk)
95% confidence intervals
methanogenesis from acetate
superpathway of guanosine nucleotides de novo bios...
pyrimidine deoxyribonucleotide phosphorylation
"superpathway of glycerol degradation to 1,3-propa...
superpathway of guanosine nucleotides de novo bios...
pyrimidine deoxyribonucleotides de novo biosynthes...
N10-formyl-tetrahydrofolate biosynthesis
lactose and galactose degradation I
superpathway of pyrimidine ribonucleotides de novo...
dTDP-L-rhamnose biosynthesis I
superpathway of pyrimidine deoxyribonucleotides de...
"superpathway of histidine, purine, and pyrimidine...
superpathway of purine nucleotides de novo biosynt...
pyrimidine deoxyribonucleosides salvage
L-arginine biosynthesis II (acetyl cycle)
tetrapyrrole biosynthesis II (from glycine)
tetrapyrrole biosynthesis I (from glutamate)
L-arginine biosynthesis I (via L-ornithine)
glycogen degradation I (bacterial)
L-arginine biosynthesis IV (archaebacteria)
superpathway of purine nucleotides de novo biosynt...
superpathway of pyrimidine deoxyribonucleotides de...
pyrimidine deoxyribonucleotides de novo biosynthes...
starch degradation V
tRNA processing
"arginine, ornithine and proline interconversion"
adenine and adenosine salvage III
androstenedione degradation
superpathway of polyamine biosynthesis II
CDP-diacylglycerol biosynthesis I
CDP-diacylglycerol biosynthesis II
adenosine deoxyribonucleotides de novo biosynthesi...
guanosine deoxyribonucleotides de novo biosynthesi...
superpathway of β-D-glucuronide and D-glucuro...
reductive acetyl coenzyme A pathway
O-antigen building blocks biosynthesis (E. coli)
heme biosynthesis II (anaerobic)
L-lysine biosynthesis I
L-tryptophan biosynthesis
homolactic fermentation
superpathway of hexuronide and hexuronate degradation
D-fructuronate degradation
D-galacturonate degradation I
urate biosynthesis/inosine 5'-phosphate degradation
1.24e-3
1.70e-3
1.92e-3
2.46e-3
2.58e-3
3.77e-3
4.10e-3
5.27e-3
5.48e-3
7.34e-3
7.59e-3
8.35e-3
8.74e-3
9.27e-3
9.45e-3
0.011
0.012
0.013
0.015
0.015
0.017
0.018
0.019
0.020
0.021
0.022
0.027
0.028
0.028
0.033
0.033
0.034
0.034
0.036
0.037
0.038
0.038
0.039
0.040
0.041
0.046
0.049
0.049
0.050
p-value (corrected)
0.0
1.3
−1.0
−0.5
0.0
0.5
1.0
Mean proportion (%)
Difference in mean proportions (%)
B
Male (12-Wk)
Female (12-Wk)
Axis 2 (27.8%)
Axis 1 (40.0%)
Male (12-Wk)
Female (12-Wk)
95% confidence intervals
Acidobacteria
Synergistetes
Cyanobacteria/Chloroplast
Poribacteria
candidate_division_WPSUnclassified1
5.19e-3
0.015
0.026
0.030
0.040
p-value (corrected)
0.0
0.0
−20
−10
0
10
20
Mean proportion (%)
Difference in mean proportions (%)
Alpha-diversity index: Chao1
Alpha-diversity index: Observed
NZM2328-Male
NZM2328-Female
Male (12-Wk)
Female (12-Wk)
95% confidence intervals
parabacteroides chongii
duncaniella muris
lactobacillus pontis
alistipes inops
bacteroides thetaiotaomicron
ligilactobacillus animalis
marinilabilia salmonicolor
monoglobus pectinilyticus
duncaniella freteri
lactobacillus intestinalis
collinsella vaginalis
2.52e-6
1.71e-3
8.72e-3
0.016
0.021
0.021
0.029
0.032
0.033
0.034
0.043
p-value (corrected)
0.0
10.5
−20
−15
−10
−5
0
5
10
15
20
Mean proportion (%)
Difference in mean proportions (%)

**Fig. S5**

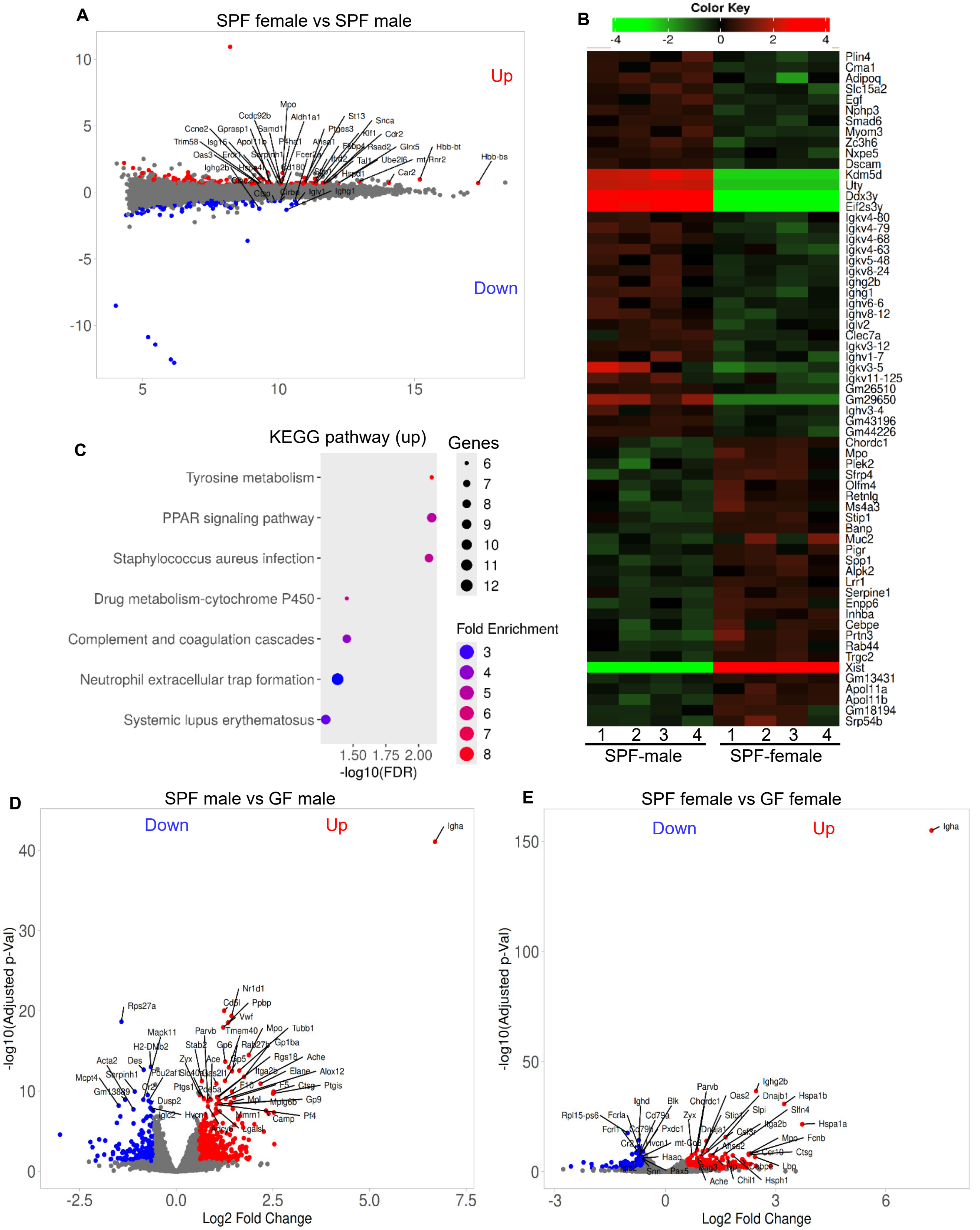


*Gene expression profiles of spleens of SPF and GF male and female NZM2328 mice.* RNA prepared from the spleen tissues of 8-week-old male and female littermates of SPF and GF NZM2328 mice were subjected to bulk-RNAseq and analyzed as described for Fig. 2. **A)** MA plot showing differentially expressed genes (DEGs; identified employing DESeq2 with a minimum fold change of 1.5-fold and FDR cutoff as 0.05) in SPF females compared to SPF males. **B)** Heatmap showing top differentially expressed genes in SPF females and SPF males. **C)** GO enrichment analysis of significantly up- regulated pathways in SPF females as compared to SPF males is shown. **D&E)** Volcano plot showing significantly up- and down- regulated genes among DEGs in SPF females and males compared to GF males and females respectively with top genes labeled.

Fig. S6A

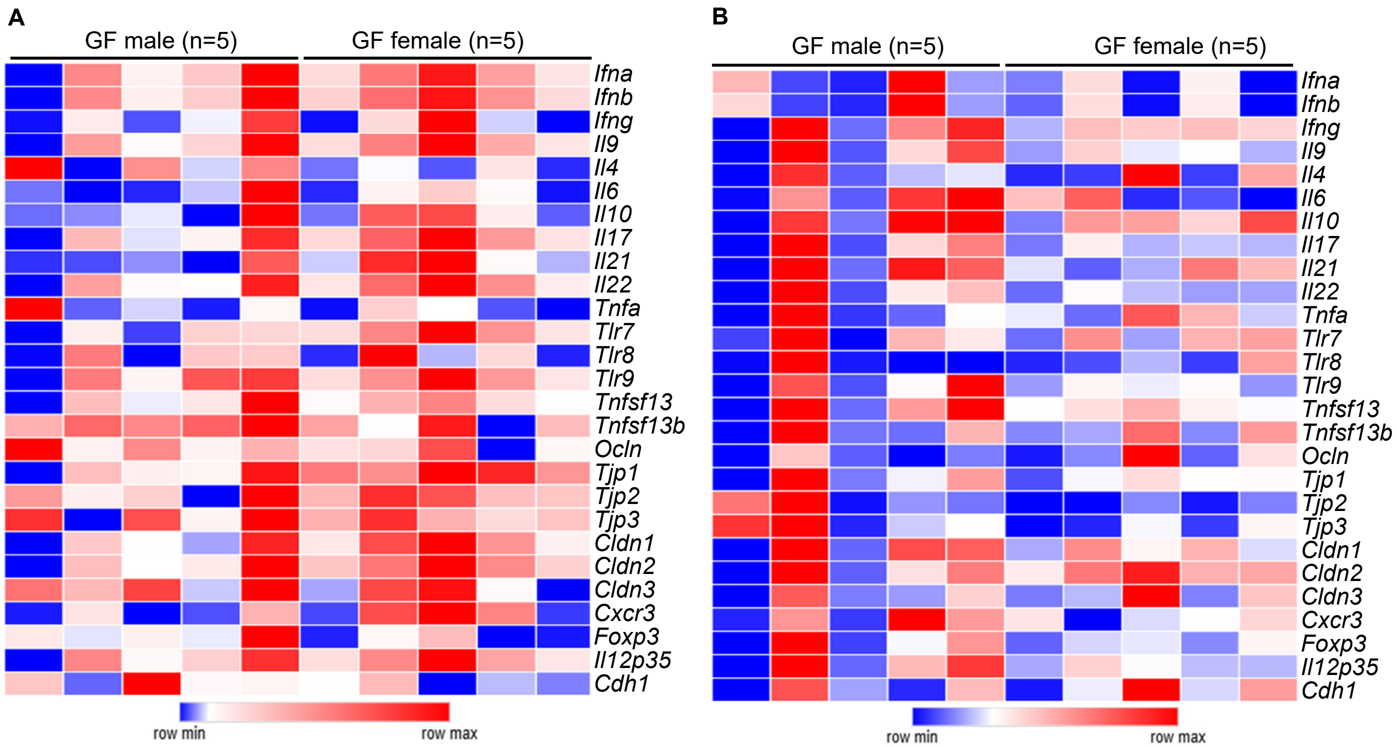


***Gene expression in intestinal tissues of GF NZM2328 mice.*** cDNA preparations from the distal colon **(A)** and distal ileum **(B)** of 16-week-old mice were subjected to qPCR to assess the expression levels of key immune and tight-junction function proteins. Expression levels of individual factors were calculated against beta-actin values and used for generating the heat-map.

**Fig. S6B**

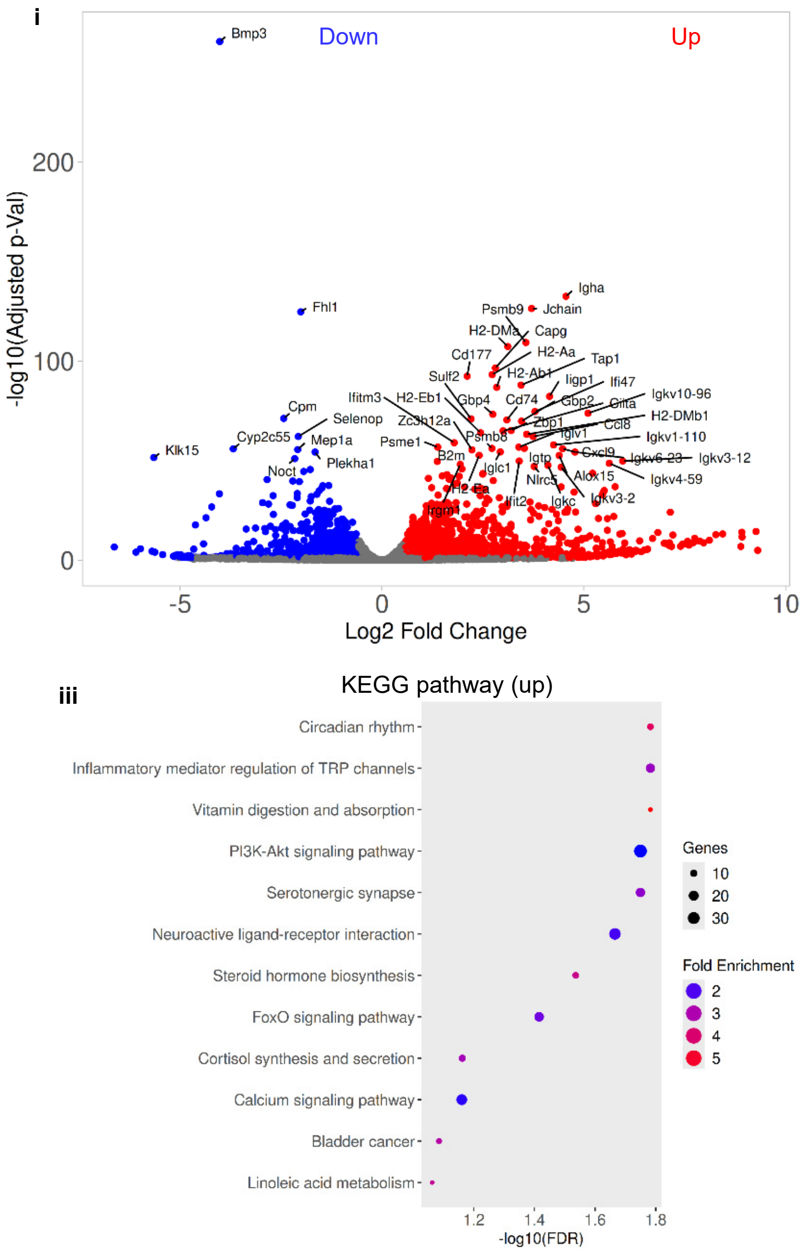


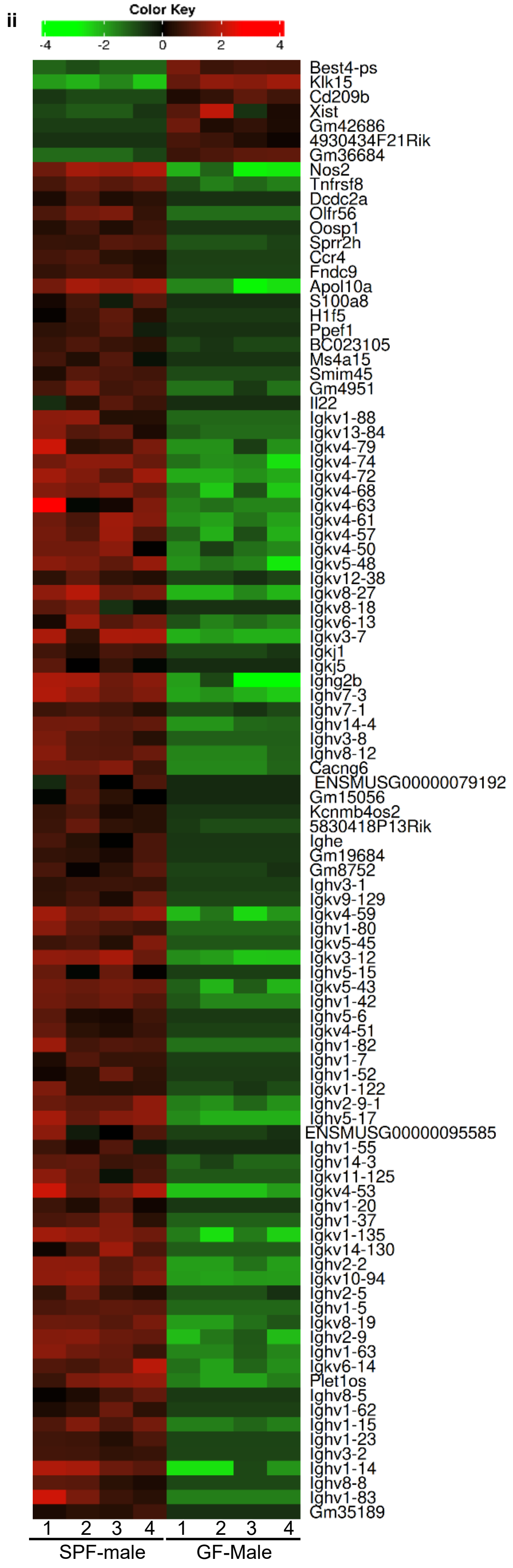


*Gene expression profiles of spleens of SPF and GF male NZM2328 mice.* RNA prepared from the distal colon tissues of 8-week-old male SPF and GF NZM2328 mice were subjected to bulk-RNAseq and analyzed as described for Fig. 2. **i)** Volcano plot showing differentially expressed genes (DEGs; identified employing DESeq2 with a minimum fold change of 1.5-fold and FDR cutoff as 0.05) in SPF males compared to GF males. **ii)** Heatmap showing top differentially expressed genes in SPF males and GF males. **iii)** KEGG enrichment analysis of significantly up-regulated pathways in SPF males as compared to GF males is shown.

### Fig. S6C

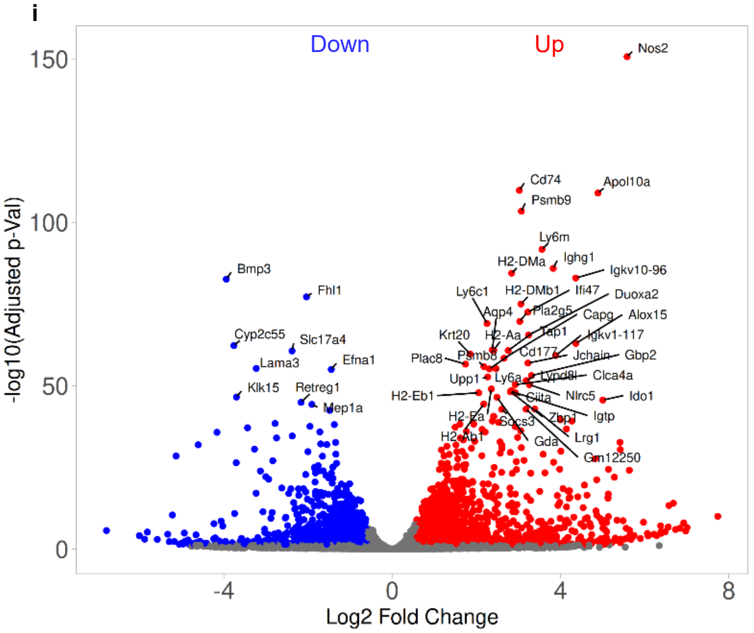


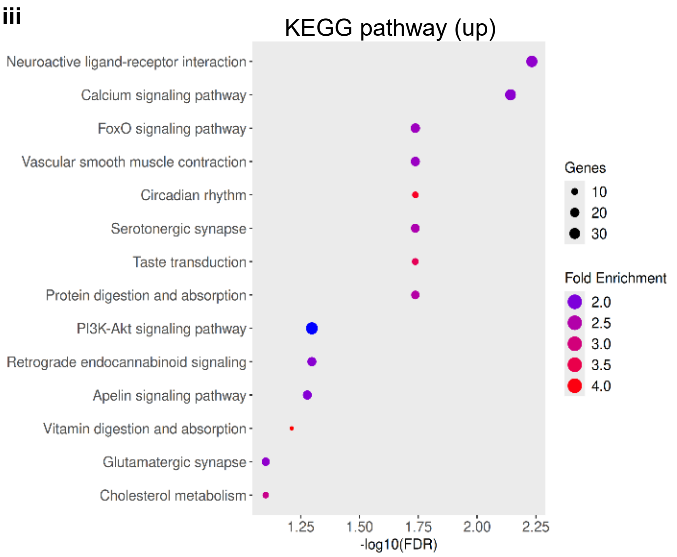


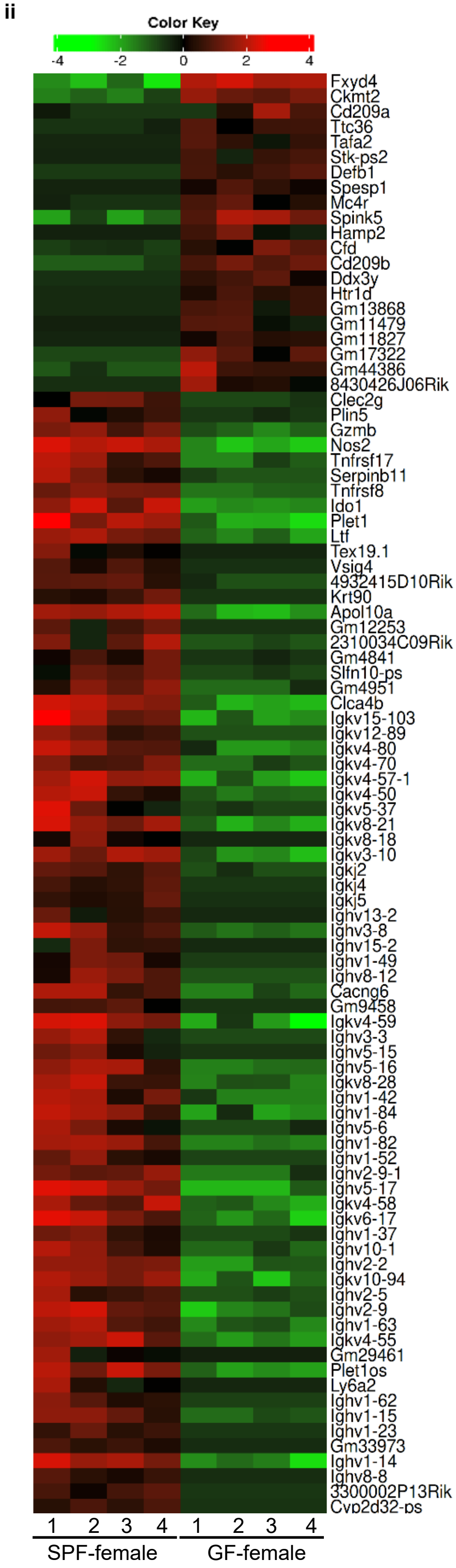


*Gene expression profiles of spleens of SPF and GF female NZM2328 mice.* RNA prepared from the distal colon tissues of 8-week-old female SPF and GF NZM2328 mice were subjected to bulk-RNAseq and analyzed as described for Fig. 2. **i)** Volcano plot showing differentially expressed genes (DEGs; identified employing DESeq2 with a minimum fold change of 1.5-fold and FDR cutoff as 0.05) in SPF females compared to GF females. **ii)** Heatmap showing top differentially expressed genes in SPF females and GF females. **iii)** KEGG enrichment analysis of significantly up-regulated pathways in SPF females as compared to GF females is shown.
